# A human pathogenic bacterium *Shigella* proliferates in the plant through the adoption of type III effectors for Shigellosis

**DOI:** 10.1101/292201

**Authors:** Sung Hee Jo, Jiyoung Lee, Eunsook Park, Dong Wook Kim, Dae Hee Lee, Choong Min Ryu, Doil Choi, Jeong Mee Park

**Affiliations:** Plant Systems Engineering Research Center, Korea Research Institute of Bioscience & Biotechnology (KRIBB), Daejeon, South Korea; Department of Biosystems and Bioengineering, KRIBB School of Biotechnology, Korea University of Science and Technology (UST), Daejeon, South Korea; Biological Resource Center, KRIBB, Jeongeup, South Korea; Plant Immunity Research Center, Department of Plant Science, College of Agriculture and Life Sciences, Seoul National University, Seoul, South Korea; Department of Pharmacy, College of Pharmacy, Hanyang University, Ansan, South Korea; Institute of Pharmacological Research, Hanyang University, Ansan, South Korea; Synthetic Biology and Bioengineering Research Center, KRIBB, Daejeon, South Korea; Infectious Disease Research Center, KRIBB, Daejeon, South Korea

**Keywords:** Alternative host, Arabidopsis plants, Enteropathogenic bacteria, PAMP-triggered immunity, Proliferation, *Shigella* spp., Split GFP, Trans-kingdom pathogenesis, Type III secretion system (T3SS), T3S effectors

## Abstract

Human enteropathogenic bacteria has been reported to be transmitted by fresh vegitables. *Shigella*, which infects primates, is reportedly transmitted by fresh vegetables; however, its molecular interactions with plants have not been elucidated. Here, we show that four *Shigella* strains, *S. boydii* (*S. b*), *S. sonnei*, *S. flexneri* 2a (*S. f* 2a), and *S. flexneri* 5a (*S. f* 5a), proliferated at different levels in *Arabidopsis thaliana*. Microscopic studies revealed that these bacteria were present inside leaves and damaged plant cells. Green fluorescent protein (GFP)-tagged *S. b* and *S. f* 5a colonized in leaves only, and *S. f* 2a colonized both leaves and roots. Using mutants lacking type III secretion systems (T3SS), we found that T3SS of *Shigella* that regulate the pathogenesis of Shigellosis in humans also play a central role in proliferation in *Arabidopsis.* Strikingly, the immunosuppressive activity of two T3S effectors, OspF and OspG, were needed for the proliferation of *Shigella* in *Arabidopsis*. Of note, delivery of OspF or OspG effectors inside of plant cells upon *Shigella* inoculation was confirmed by using a split GFP system, respectively. These findings demonstrate that the human pathogen *Shigella* can proliferate in plants by adoption of immunosuppressive machinery for its original host human.

## Introduction

Human hosts are primarily infected by the ingestion of water or food contaminated with bacteria from feces (the fecal-oral route), including ingestion of undercooked contaminated meats and improperly washed contaminated vegetables (Mead et al., 1999; Weir, 2002; Gupta et al., 2004). Fresh fruits and vegetables, such as lettuce, tomatoes, and green peppers, are responsible for the widespread transmission of food-borne infections by *Salmonella* or *Shigella* (Guchi and Ashenafi, 2010; Semenov et al., 2010; Gu et al., 2013). These observations suggest that human pathogenic bacteria use plants as alternative hosts as a stable environmental niche. Several human pathogenic bacteria, including *Salmonella enterica* serovar Typhimurium, *Escherichia coli* O157:H7, and *Pseudomonas aeruginosa*, are known to use plants as alternative hosts (Plotnikova et al., 2000; Semenov et al., 2010). These bacteria can attach to the plant surface, and actively invade and proliferate in plant tissues (Cevallos-Cevallos et al., 2012; Martínez-Vaz et al., 2014). In particular, several enteropathogenic bacteria, including *E. coli* O157:H7 and *Salmonella*, spread within plants through vascular tissues after infection via contaminated water (Solomon et al., 2002).

*Shigella* is a human-adapted pathogen that infects the host via multiple transmission routes. It is a non-motile, rod-shaped, facultative intracellular and invasive pathogen, very closely related to *Escherichia coli*. Based on the carbohydrate composition of the O-antigen, i.e., the polysaccharide component of the lipopolysaccharide molecule that is the major bacterial surface antigen, *Shigella* is classified into four serogroups. These have been given species designations, namely, *S. dysenteriae* 1 (serogroup A), *S. flexneri* (serogroup B), *S. boydii* (serogroup C), and *S. sonnei* (serogroup D) (Lindberg et al., 1991; Schroeder and Hilbi, 2008). *Shigella* spp. are important epidemic pathogens and serious public health concern in developed and developing countries. About 164,300 deaths in all age groups, and 54,900 deaths in children younger than 5 years, were reported globally in 2015 (Mortality and Causes of Death, 2016). However, the actual number of infections might be higher because mild symptoms are not reported (Mortality and Causes of Death, 2016), suggesting that the microorganism may employ a variety of survival strategies not only for human intestinal infection but also for survival in a non-adapted host. Although *Shigella* contamination has also been reported in plants (Naimi et al., 2003; Ohadi et al., 2013), it is not yet known whether the bacterium actively invades and/or proliferates inside the plant.

Unlike animals, plants have no adaptive immune system; instead, each cell possesses an innate immune system. The innate immune system in plants and animals recognizes and suppresses pathogens and has common features that are preserved throughout evolution. Plant pattern recognition receptors recognize conserved microbial or pathogen-associated molecular patterns; the pattern-triggered immunity (PTI) is activated via the mitogen-activated protein kinase (MAPK) cascades (Jones and Dangl, 2006). To suppress PTI, bacteria inject effector proteins into plant cells using type III secretion systems (T3SS). To counteract this PTI evasion response, the plant nucleotide binding-leucine rich repeat proteins recognize the pathogen effectors; effector-triggered immunity is then activated to accompany the hypersensitive response (Jones and Dangl, 2006). For human or animal intestinal bacteria to infect plants, the PTI must first be disabled. *S. enterica* serovar Typhimurium, similar to its activity in the mammalian host, uses T3SS to suppress plant immune responses (Schikora et al., 2011; Schikora et al., 2012). In particular, one of the T3S effector proteins of *S. enterica,* SpvC targets the MAPK signaling system as in animals to suppress the host PTI, when expressed in plants under the control of plant binary vector (Neumann et al., 2014).

Here, we examined the ability of four *Shigella* strains (*S. s* (Holt et al., 2012), *S. b* and *S. f* 2a (Wei et al., 2003), and *S. f* 5a (Onodera et al., 2012)) to proliferate in *Arabidopsis* plants. We found that the four strains invaded and proliferated differently in plant tissues. A Proliferation of mutants lacking the T3S effectors, i.e., noninvasive human strains, was reduced *in planta*. Reverse genetics and molecular biology experiments demonstrated that the immunosuppressive function of *Shigella* T3S effectors OspF and OspG was essential for *Shigella* proliferation in plants. Notably, we first observed the delivery of *Shigella* Type III effector proteins inside of plant cells by *Shigella* inoculation using the split GFP technology. These observations indicate that *Arabidopsis* may be useful as a model host for studying the pathogenesis of *Shigella*.

## Materials and Methods

### Plant materials and growth

*Arabidopsis thaliana* accession Columbia (*Col-0*) was used for *Shigella* infection. Briefly, *Arabidopsis* seeds were surface-sterilized in 70% (v/v) ethanol for 2 min followed by an incubation in 50% household bleach for 10 min. Seeds were washed extensively with sterile deionized water before placed on 1/2 Murashige and Skoog (MS) medium (Duchefa Biochemie, Haarlem, Netherlands) supplemented with 1% sucrose and solidified with 0.6% (w/v) agar (Murashige and Skoog, 1962). *Nicotiana benthamiana* plants were grown at 22 ± 3°C under a 16 h light/8 h dark cycle in plastic pots containing steam-sterilized mixed soil (2:1:1, v/v/v, soil/vermiculite/perlite) (Moon et al., 2016). To measure the plant immune response in terms of MAPK activity, 1 μM flg22 peptide (Cat No: FLG22-P-1; Alpha Diagnostics, Inc., San Antonio, TX, USA) was used as a positive control (Bethke et al., 2009).

### Bacterial strains, growth conditions, and plasmids

The bacterial strains and plasmids used in the study are described in Table S1. *Shigella* and *Pseudomonas* strains harboring a plasmid pDSK-GFPuv were generated by electroporation, as described previously (Wang et al., 2007; Hong et al., 2016).

*Shigella* spp. were grown at 37°C in Luria-Bertani (LB) medium or tryptic soy agar containing 0.003% (w/v) Congo red dye (Sigma-Aldrich, St. Louis, MO, USA) (Runyen-Janecky and Payne, 2002). *Pseudomonas syringae* strains were grown at 28°C (with shaking at 200 rpm) in King’s B liquid medium (Sigma-Aldrich) containing appropriate antibiotics (King et al., 1954). As a non-pathogenic control, *Escherichia coli* DH5α was grown in LB medium at 37°C with shaking (Kennedy, 1971). *Agrobacterium tumefaciens* GV2260 was grown at 28°C in LB broth with shaking at 200 rpm (Shamloul et al., 2014).

The coding region of *ospF* or *ospG* was PCR-amplified using *attB*-containing PCR primers (Table S2). The PCR fragments were cloned into the pDONR™207 vector by BP recombination using the Gateway^®^ BP Clonase™ II kit (Invitrogen, Carlsbad, CA, USA). The products were then transferred to pBAV178 (for AvrRpt2 fusion) or pBAV179 (for HA fusion) vectors by LR recombination (Gateway^®^ LR Clonase™ II, Invitrogen). pBAV178, pBAV179, and pME6012 (empty vector control) were kindly provided by Dr. Jean T. Greenberg (University of Chicago) (Vinatzer et al., 2005).

### Bacterial inoculation assay *in planta*

*Arabidopsis* seedlings (2 weeks old) grown in 1/2 MS medium were used for flood-inoculation (Ishiga et al., 2011). Briefly, 10 *Arabidopsis* seedlings in one container were incubated for 3 min with 35 ml of each bacterial strain suspension (5 × 10^6^ or 5 × 10^5^ cfu/ml) containing 0.02% Silwet L-77 (Lehle Seeds, Round Rock, TX, USA) or buffer. After the bacterial suspensions were removed by decantation, plates containing inoculated plants were incubated in a growth room (23 ± 2°C, 16 h light/8 h dark). Bacterial cell counts from inoculated plants were monitored as described previously (Ishiga et al., 2011). Three inoculated seedlings in one container were harvested by cutting the hypocotyls, and total fresh weight was measured. The cfu were normalized to cfu/mg using sample weight. The cfu of seedlings in three separate containers (as biological replicates) were measured. In addition, the bacterial population was evaluated in more than three independent experiments conducted successively under the same conditions.

To assess root invasion, 10-d-old *Arabidopsis* seedlings grown vertically in 1/2 MS medium were inoculated by dropping 2.0 μl of bacterial suspension (5 × 10^7^ cfu/ml) onto the root tips. Symptoms were observed under white light, and bacterial proliferation was monitored at 5 dpi by observation of GFP-expressing bacteria under UV light. Three biological replicates were generated in separate plates, and three independent experiments were conducted under the same conditions.

To analyze the effector secretion, pBAV178 was used to transform *P. syringae* pv. *tomato* DC3000 (*Pst*) by electroporation (Cadoret et al., 2014). Bacterial suspensions (2.5 × 10^8^ cfu/ml) were used to syringe-infiltrate *Arabidopsis* leaves; 2 d after the infiltration, the hypersensitive response was assessed by trypan blue staining (Koch and Slusarenko, 1990).

To test the virulence of *Shigella* effectors via *Pst,* plasmid pBAV179 was used to transform *Pst*; 5 × 10^7^ cfu/ml bacterial suspensions containing 0.02% (w/v) Silwet L-77 were used to spray-infect *Arabidopsis* leaves. After infection, the plants were covered with a clear lid to maintain humidity and transferred to a growth room (22 ± 3 °C, 16 h light/8 h dark). The symptoms and bacterial proliferation were assessed at 4 dpi.

### *Shigella* infection to visualize of effector secretion into plant cells

Transgenic *Arabidopsis* expressing Nucleus-targeted Nu-sfGFP1-10^OPT^ were flood inoculated (5 × 10^5^ cfu/ml) with *S. f* 5a, *S. f 5a*::OspF-sfGFP11 or *S. f 5a*::OspG-sfGFP11. To produce the sfGFP11-fused *Shigella* effectors, *OspF* or *OspG* was transferred to pEP119T by LR recombination. At specific time points, *Arabidopsis* leaf discs were observed under a confocal microscope (Park et al., 2017). Each experiment included at least three independent plants.

### Microscopy

For SEM, flood-inoculated *Arabidopsis* leaves were fixed in 4% (w/v) paraformaldehyde and dehydrated in an ethanol series (30%, 50%, 70%, 96%, and 100%). The fixed leaves were then dried, coated with gold-palladium, and visualized using a scanning electron microscope (LEO 1455VP, Oberkochen, Germany) (Plotnikova et al., 2000). For TEM, flood-inoculated *Arabidopsis* leaves were cut off, fixed overnight in 2.5% (w/v) glutaraldehyde, post-fixed in 2% (w/v) osmium tetroxide, dehydrated in ethanol, and embedded in the resin. After staining in 2% (w/v) uranyl acetate and lead citrate, samples were observed under an electron microscope (Bio-TEM; Tecnai G2 Spirit Twin; FEI, USA) (Chae and An, 2016).

For fluorescence confocal microscopy, expression of GFP-labeled bacteria or GFP-tagged *Shigella* effector proteins in plants was observed under a Nikon laser scanning confocal microscope C2 (Nikon, Tokyo, Japan) using filter sets for GFP (λ_ex_, 488 nm; λ_em_, 505–530 nm) or RFP (λ_ex_, 561 nm; λ_em_, 570–620 nm). For each microscopic method, three leaves were used per treatment and at least three microscopic fields were observed for each leaf, including the control.

### Expression of *Shigella* virulence genes in *Arabidopsis* plants

Total RNA was extracted from *Shigella*-infected leaves (from three plants) using RNAiso plus (Cat No: 9108; TaKaRa, Otsu, Japan), according to the manufacturer’s protocol. RT-PCR was performed using M-MLV reverse transcriptase (Invitrogen), according to the manufacturer’s instructions. Quantitative RT-PCR was carried out in a CFX Connect™ Real Time System (BioRad, Hercules, CA, USA) using iQ™ SYBR^®^ Green Supermix (BioRad) and primers specific for target genes (*ipaB*, *ipaC*, *icsA*, *icsB*, *virB*, and *virF*; Table S1 (Bando et al., 2010)). The qRT-PCR results were normalized to the expression of 16s rRNA.

### Immunoblotting

Total protein was extracted from *Shigella-* or *Agrobacterium-*infected leaves (from three plants) using denaturing extraction buffer [150 mM NaCl, 50 mM Tris-HCl (pH 7.5), 5 mM EDTA, 0.1% Triton-X, 1× protease inhibitor cocktail (Roche, Basel, Switzerland), 0.4 M DTT, 1 M NaF, and 1 M Na_3_VO_3_]. The extracted proteins were separated on 12% SDS-PAGE gels and transferred to a PVDF membrane (Pierce, Rockford, IL, USA). Antibodies specific for phospho-p44/p42 MAPK ERK1/2 (Cat No: 4377; Cell Signaling Technology, Danvers, MA, USA) and ERK1 (Cat No: sc-94; Santa Cruz, Dallas, TX, USA) were used for immunoblot analyses. Target proteins were detected using ECL plus reagent (GE Healthcare, Wauwatosa, WI, USA) and visualized using an Alliance 9.7 chemiluminescence imaging system (UVITEC, Cambridge, UK).

### Statistical analysis

All data are expressed as the mean ± SD. The statistical significance of bacterial cell growth in infected plants was examined using Student’s t-test (Microsoft Office Excel) and ANOVA (SPSS v.18; IBM, Armonk, NY, USA) (Moon et al., 2016). Asterisks and letters indicate significant differences between samples (*P* < 0.05).

## Results

### Four *Shigella* spp. strains interact differently with *Arabidopsis*

To observe the behavior of the human pathogen *Shigella* in plants, we investigated the interaction of four *Shigella* spp. strains representing three serogroups (*S. b*, *S. s*, *S. f* 2a, and *S. f* 5a), with the model plant *A. thaliana*. *Shigella* is a water-borne pathogen, with infection spreading via contaminated water (Pandey et al., 2014). Hence, we chose to use a flood-inoculation approach (Ishiga et al., 2011), which is thought to mimic natural inoculation. Infection of the phytopathogen *Pseudomonas syringae* pv. *tomato* DC3000 (*Pst*) was observed as a positive control and *Pst* Δ*hrcC*, a mutant lacking the T3SS of *Pst*, and the non-pathogenic bacterium *E. coli* DH5α were used as negative controls for infection. When 2-week-old *Arabidopsis* seedlings were inoculated with *S. s* or *S. f* 2a, clear infections were observed, such as yellowing and necrosis of the leaves, while mild symptoms were detected after inoculation with *S. b* and *S. f* 5a (Figure 1A) (Liu et al., 2015).

**Figure 1.**
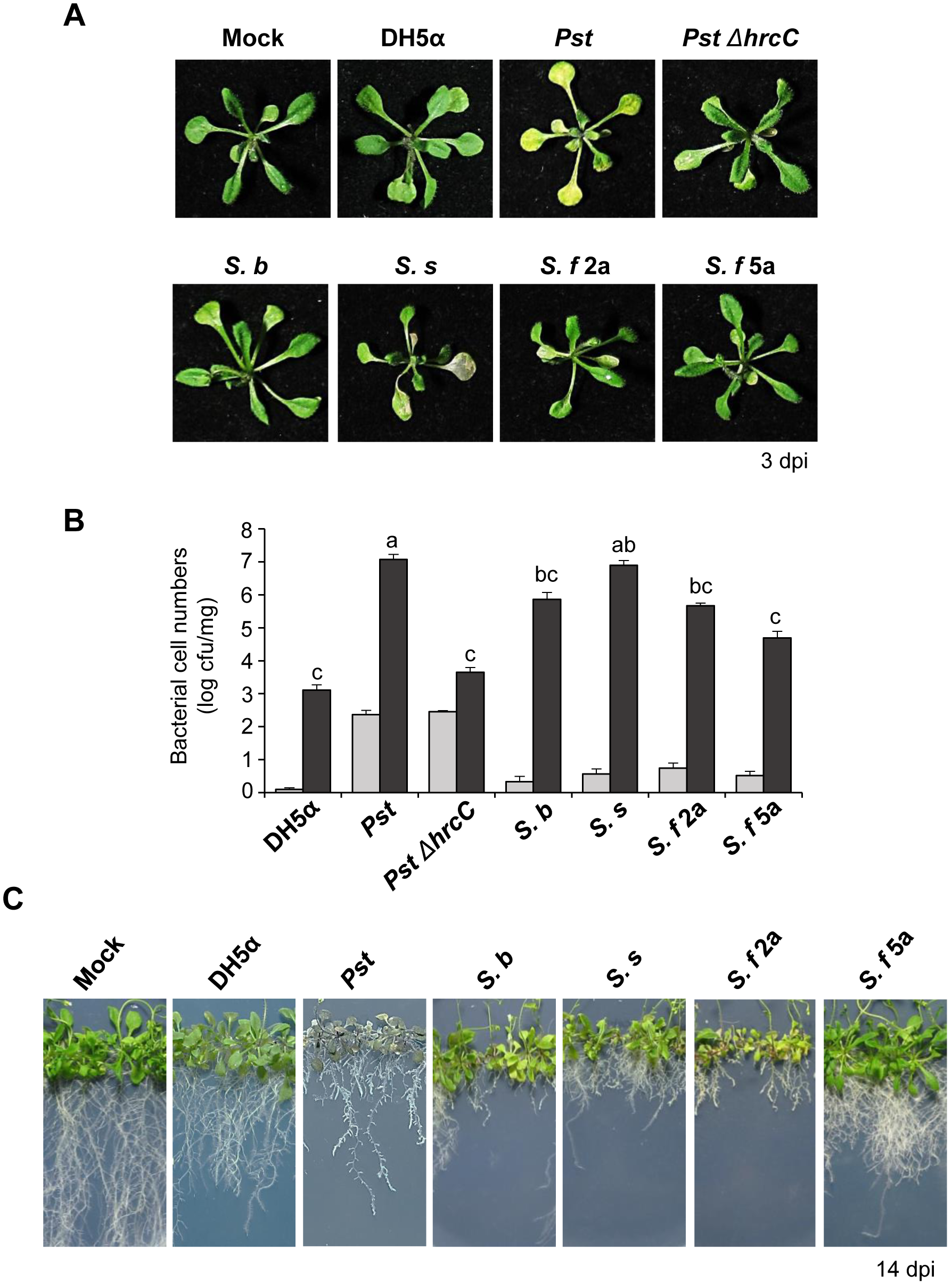
*Shigella* proliferates and induces disease symptoms in *Arabidopsis* plants. (A, B) *Arabidopsis* seedlings in 1/2 MS medium were flood-inoculated with distilled H_2_O containing 0.025% Silwet L-77 (Mock) or bacterial suspensions (5 × 10^5^ cfu/ml). (A) The symptoms of *Arabidopsis* inoculated with *Shigella* spp. by flood-inoculation. Photographs of representative symptoms were taken at 3 dpi. (B) Bacterial cell numbers were evaluated on Days 0 (gray bar) and 3 (black bar) after flood-inoculation. Plants were surface-sterilized with 5% H_2_O_2_ and washed several times with sterile distilled water to remove unattached bacteria from their surface, before evaluating attached and proliferated bacterial population. The bars represent the mean ± SD of three replicates and the different letters indicate significant differences between samples (P < 0.05, one-way ANOVA). (C) *Arabidopsis* seedlings in 1/2 MS medium were infected with 2 μl of buffer or bacterial suspensions (5 ×10^7^ cfu/ml) by dropping onto the root tips. The photographs were taken 14 dpi and are representative of three independent experiment. All experiments were repeated three times independently and representative results are shown.

In addition to observing the symptoms, the bacterial growth *in planta* was also evaluated to detect initial plant adherence and proliferation. Early attachment of all *Shigella* strains and DH5α was more than 10 times lower than that of *Pst* and *Pst* Δ*hrcC* (Figure 1B; Day 0, grey bars). At 3 days post-inoculation (dpi), the cell number of all strains significantly increased than initial bacterial cells, although the extent of cell amplification differed depending on the *Shigella* strain (Figure 1B; black bars). Notably, *S. s* cell numbers increased over 10^5^ times, comparable to the proliferation of *Pst*. By contrast, the interaction between *S. f* 5a and *Arabidopsis* was similar to that between the plant and non-pathogenic DH5α.

Previous studies demonstrate that several human pathogenic bacteria, including *E. coli* O157:H7 and *Enterococcus faecalis*, invade the leaves and roots of *A. thaliana* (Jha et al., 2005; Deering et al., 2012). Therefore, to determine whether *Shigella* strains invade through *A. thaliana* roots, we dropped bacterial solutions onto root tips and observed symptoms in *Arabidopsis* plants for 14 d. Interestingly, all the strains inhibited Arabidopsis root growth (Figure 1C). Interestingly, *S. b* strain caused severe root growth inhibition unlike the mild disease symptom of the leaves after the infection of this strain (compare Figure 1A, 1C). Inoculation of *S. f 5a* caused slight inhibition of root growth but caused much less damage to the plant than other strains. Taken together, these results indicated that, although the proliferative capacity of the *Shigella* strains differs, the cells can invade and colonize the apoplast of plants, thereby causing structural damage to the host.

### Penetration and subsequent internalization of *Shigella* spp. into *Arabidopsis*

Since we observed that *Shigella* proliferate and induce disease-like symptoms in *Arabidopsis*, we examined whether the bacterium multiplies on the leaf surface and in the intercellular space (apoplast) by scanning electron microscopy (SEM) and transmission electron microscopy (TEM), respectively. *Pst*, known to infects plants via open stomata (Panchal et al., 2016), colonized guard cells at 24 h post-infection (Figure 2A). Similarly, all tested *Shigella* strains clustered around guard cells and the surface of epidermal cells. *S. s* and *S. b* formed relatively broad clusters in the surrounding areas, including guard cells (Figure 2A). In particular, *S. b* and *S. f* 2a intensely colonized guard cells (Figure 2A), leading us to speculate that they enter plants via open stomata, similar to *Pst*.

**Figure 2.**
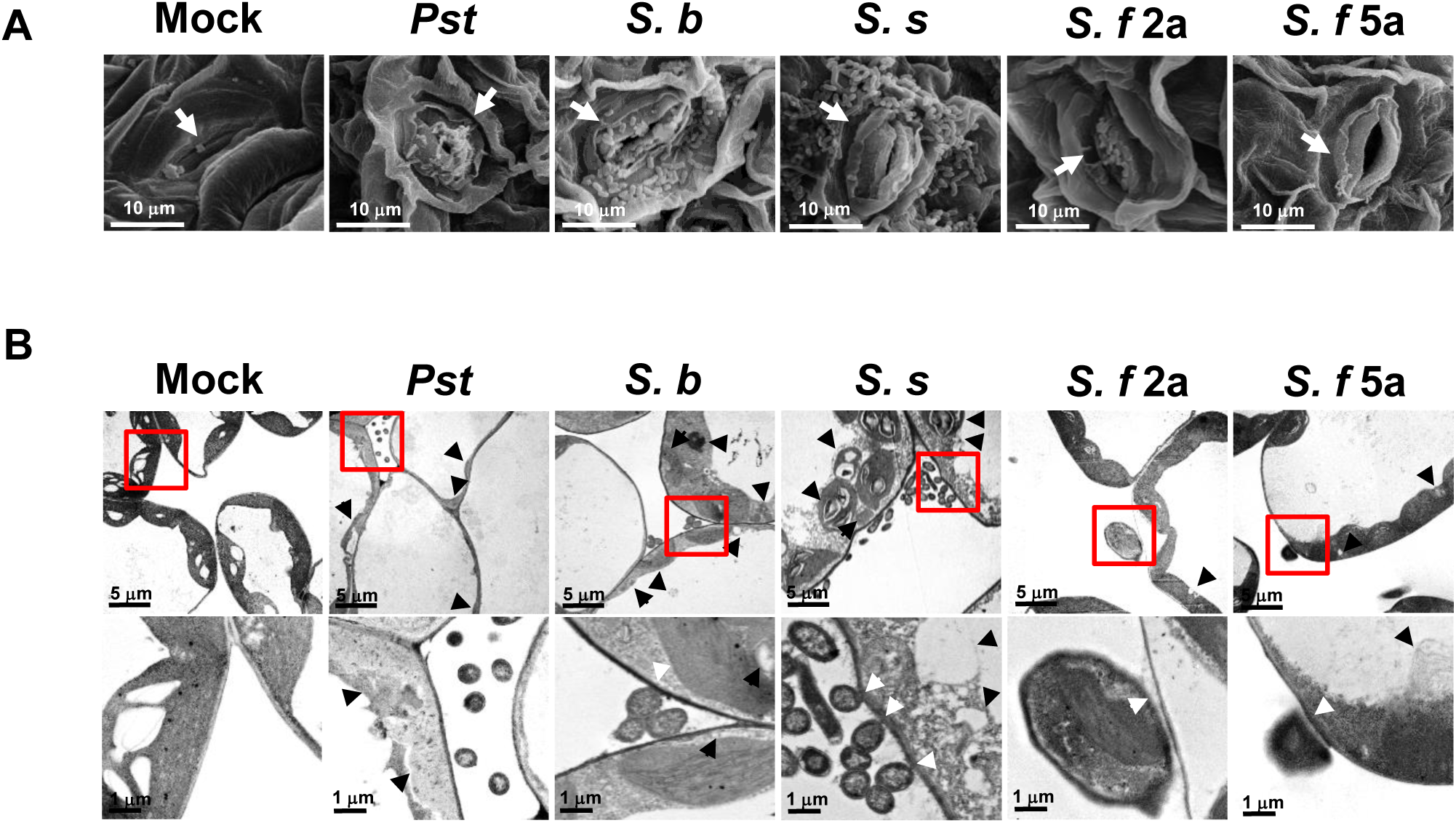
*Shigella* clusters around guard cells and localizes in the apoplast affecting plant cell structures. (A, B) *Arabidopsis* seedlings were flood-inoculated with a buffer or bacterial suspensions (5 × 10^6^ cfu/ml). (A) After 24 h, the leaves were fixed and analyzed under a SEM. The *Pst* cells were observed over the stomata. *Shigella* strains were associated with the stomata. Bar, 10 μm. Representative image showing bacteria around guard cells (indicated by white arrows). (B) After 24 h, the leaves were evaluated under a TEM. The *Shigella* strains colonized the intercellular spaces. Red boxed regions in the micrographs at the upper panel were maginified at the bottom panel. Bar, 5 μm (upper panels). Bar, 1 μm (bottom panels). TEM images revealed the *Shigella* (white arrowheads in the bottom panel) attached to the cell wall in the intercellular spaces and altered mesophyll cells. Black arrowheads at the upper panel indicate separation of the plasma membrane from the cell wall, the abnormal organelles and disruption of chloroplasts. The results are representative of two independent experiments.

Most plant pathogenic bacteria infect and colonize the apoplast (Sattelmacher, 2009; Gao et al., 2016). Therefore, we investigated whether *Shigella* is capable of intercellular colonization and causing damage to plant cells. Indeed, the TEM images revealed colonization of *Shigella* in the intercellular spaces and attachment to the host cell walls (Figure 2B). The presence of the microbes in the intercellular space resulted in the alteration of the host organelle structure, such as the separation of the plasma membrane from the cell wall, the liberation of cell organelles, and disruption of chloroplasts (Figure 2B (Gao et al., 2016)). This effect was most pronounced in plant cells inoculated with *S. b* and *S. s*; further, *S. b* and *S. s* were more commonly found in the intercellular spaces than *S. f* 2a and *S. f* 5a.

To observe internalization of bacterial cells in the plant via root cells, we attempted to label the four *Shigella* strains with a green fluorescent protein (GFP) and obtained *S. b*, *S. f* 2a, and *S. f* 5a strains labeled with GFP successfully. GFP labeling did not affect the bacterial growth in plants (Figure S2). Bacterial suspensions were dropped onto the root tips of vertically grown *Arabidopsis* plants. Five days later, whole plants and root tissues were photographed under ultraviolet (UV) light to observe the distribution of fluorescently labeled bacteria (Figure 3A). In accordance with the disease phenotypes observed, *Pst* (which is a foliar pathogen) exhibited strong fluorescence in the leaves, despite the fact that it was applied to root tips (Figure 3A). To observe *Pst* in root tissues, inoculated roots were disinfected with hypochlorite solution and observed under a fluorescence microscope. GFP fluorescence was observed only in epidermal cells after application of *Pst* to root tips (Figure 3B). Consistantly, this finding indicates that *Pst* infects throughthe leaf tissues, not by invading root tissues or colonizing the roots. Interestingly, when GFP-labeled *S. f 2a* was applied to the root tip, strong green fluorescence was observed in both roots and leaves, while other GFP-labeled *Shigella* inoculated plants showed only weak fluorescence in leaves (Figure 3A). In root tissues, GFP fluorescence was observed in root endodermal cells only in *S. f* 2a-treated plants (Figure 3B). Taken together, these results indicate that *Shigella* invades plant leaves and roots in a strain-dependent manner, and then moves along the surface and through the internal vascular tissues of the plant.

**Figure 3.**
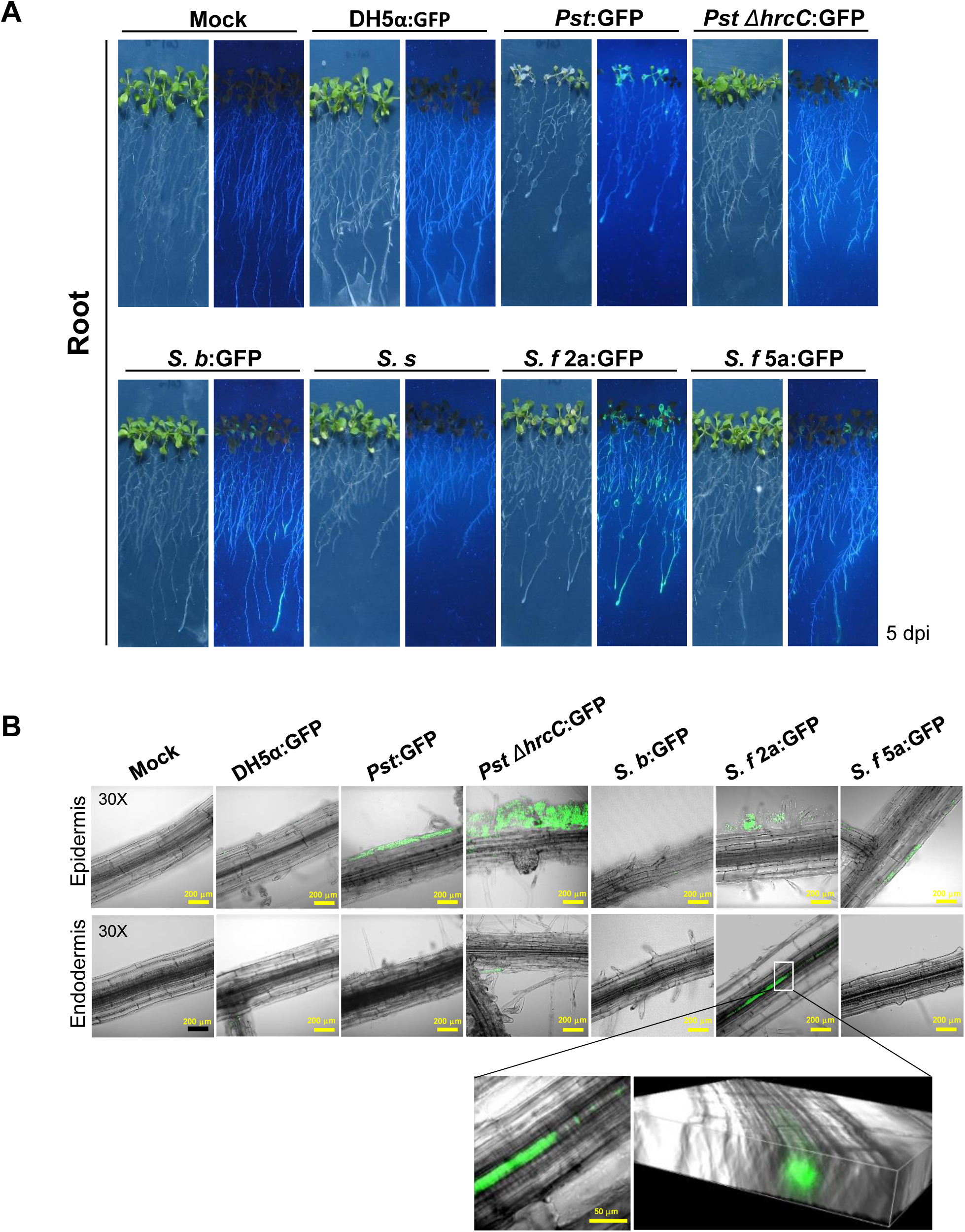
*Shigella S. f 2a* invades and colonizes plant roots. (A, B) Root tips of *Arabidopsis* seedlings were drop-inoculated with GFP-labeled bacterial suspensions (5 × 10^7^ cfu/ml). (A) Bacterial colonization of inoculated plants was photographed under white and UV light at 5 dpi. (B) GFP-labeled *Shigella* are localized in the epidermal or endodermal cells of *Arabidopsi*s roots. GFP images were taken using a confocal microscope. Higher magnification micrographs and 3D Raman confocal volume images show internalization of *S. f* 2a in *Arabidopsis* roots. All experiments were repeated at least three times, and representative results are shown.

### *Shigella* T3S effectors are necessary for attachment and multiplication in *Arabidopsis*

T3SS is the crutial determinant for the virulence of many Gram-negative bacteria, including animal and plant pathogens (He, 1998; Ogawa et al., 2008; Buttner and He, 2009). Thus, we investigated if pathogenic proteins for animal host infections are required for invasion and multiplication of *Shigella* in plants. To study the biological role of the T3SS in the interaction between *Shigella* and plants, we used noninvasive variants of *S. f* 2a and *S. f* 5a (strains Δvp and BS176, respectively) (Sansonetti et al., 1982; Wenneras et al., 2000; Shim et al., 2007). To facilitate observation of bacterial invasion in living plants, the noninvasive variants of the *Shigella* strains were also labeled with GFP, and bacterial growth and expression of effector proteins were verified (Figure S1-S3). Bacterial proliferation in plants after inoculation with Δvp or BS176 strains was 10 times lower than that after inoculation by parental *Shigella* strains; similar results were observed for GFP-labeled *Shigella* strains and mutants (Figure 4A). To examine the involvement of the T3SS in bacterial invasion of the plant surface, leaves were flood-inoculated with GFP-labeled bacteria, and leaf surfaces were observed 24 h later (Figure 4B). Examination under UV light revealed that the levels of GFP-labeled *S. f* 2a and *S. f* 5a on leaf surfaces were higher than those of Δvp and BS176, especially in open stomata (Figure 4B). We also generated a non-pathogenic mutant of *S.s* and have tested if the severe infection of *S.s* strain is T3SS dependent (Figure S4). When plants were inoculated with this T3SS-deficient *S. s* strain, disease phenotypes and bacterial growth were significantly reduced (Figure S4), as observed with Δvp and BS176. Altogether, these results suggested that the T3SS of *Shigella* that operate during infection in human is also required for the colonization in plants.

**Figure 4.**
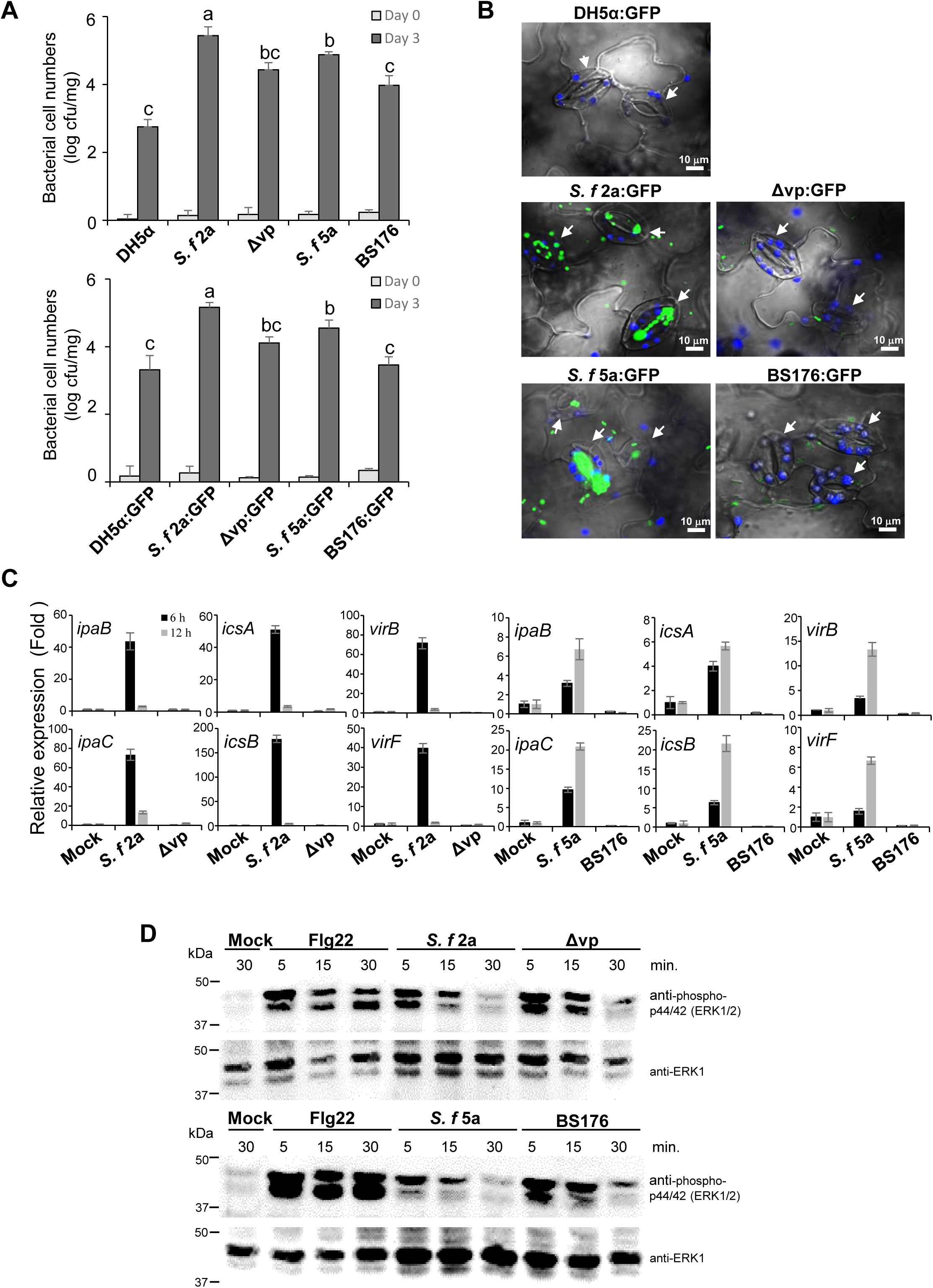
The virulence plasmid-deficient mutant of *Shigella* is impaired in the *Arabidopsis* proliferation. (A, B) *Arabidopsis* seedlings in 1/2 MS medium were flood-inoculated with *S. f* 2a, Δvp, *S. f* 5a, or BS176, and their GFP-labeled variants (5 × 10^5^ cfu/ml). (A) The bacterial cell numbers *in planta* were determined at 0 and 3 dpi. Bars represent the mean ± SD of three replicates and the different letters indicate significant differences between samples (*P* < 0.05, one-way ANOVA). (B) Images of GFP-labeled *Shigella* in leaf epidermal cells of *Arabidopsis* acquired at 24 hpi under a fluorescence confocal microscope. Bar, 10 μm. The blue represents auto-fluorescence of chlorophyll. All experiments were repeated at least three times, and representative results are shown. (C, D) *Arabidopsis* leaves were syringe-infiltrated with buffer, 1 µM flg22, or bacterial suspension (5 × 10^6^ cfu/ml), and samples were collected at the indicated times. (C) Transcription of *Shigella* effectors (*ipaB*, *ipaC*, *icsA*, *icsB*, *virB*, and *virF*) in inoculated *Arabidopsis* leaves was analyzed by qRT-PCR. qRT-PCR results were normalized to expression of 16s rRNA. Expression of effectors by plasmid-deficient mutant strains was compared with that in the WT. Data are expressed as the mean ± SD of three replicates. (D) Immunoblotting was performed using either anti-phospho-p44/42 (ERK1/2, upper panels) or anti-ERK1 (lower panels) antibodies. All experiments were repeated at least three times, each with similar results.

To further investigate involvement of *Shigella* T3S effectors in plant interactions, we measured expression of critical virulence effectors related to human pathogenesis, including *ipaB*, *ipaC*, *icsA*, *icsB*, *virB*, and *virF*, in *S. f* 2a-, *S. f* 5a-, Δvp-, and BS176-infected *Arabidopsis* plants (Bando et al., 2010). These effectors play a role in mammalian cell lysis (*ipaB*, *ipaC*), intracellular spread (*icsA*, *icsB*), and regulation of virulence factor (*virF*, *virB*) (Ogawa et al., 2008). Total RNAs were isolated from *Arabidopsis* leaves at 6 and 12 h after *Shigella* inoculation, and changes in expression of virulence genes were confirmed by quantitative RT-PCR (Figure 4C). Expression of all *Shigella* virulence genes examined in WT *S. f* 2a- or *S. f* 5a-inoculated *Arabidopsis* leaves increased. Interestingly, induction was higher and faster in *S. f* 2a-treated plants than in *S. f* 5a-treated plants, which is in agreement with the earlier results showing that *S. f* 2a was more pathogenic to plants than *S. f* 5a. Expression of virulence genes was not detected in plants inoculated with non-pathogenic mutants Δvp or BS176, similar to buffer-treated control plants (Figure 4C), suggesting that common virulence factors regulate interactions with both plants and human intestinal cells.

To investigate the plant innate immune responses to *Shigella* inoculation, MAPK phosphorylation was evaluated. The activation of MAPK by phosphorylation is a conserved response of the earliest microbe-triggered immune signaling in both plants and animals (Zipfel, 2009). The flg22 peptide is a representative microbe-associated molecular pattern in plants (Bethke et al., 2009). In plants treated with flg22, pronounced MAPK phosphorylation was apparent within 5 min of treatment and this response lasted up to 30 min (Figure 4D). On the other hand, MAPK phosphorylation in plants treated with *S. f* 2a or *S. f* 5a was reduced; from 15 min on, it was strongly suppressed and almost completely disappearing after ca. 30 min (Figure 4D). Meanwhile, in plants treated with the virulence plasmid-deficient mutants, Δvp or BS176, MAPK activation was recovered, in contrast to *S. f* 2a- or *S. f* 5a-treated plants, although the degree and duration of the phosphorylation were lower than those elicited by the flg22 treatment (Figure 4D). These observations indicated that *Shigella* suppresses the innate immunity of *Arabidopsis* via its T3SS.

### Suppression of immune signaling in *Arabidopsis* plants by *Shigella* T3S effectors OspF and OspG

*Shigella* OspF has been reported to inhibit MAPK signaling, which is conserved in plants and animals (Arbibe et al., 2007; Li et al., 2007). OspG is an essential immunosuppressive effector protein secreted at the later stages of infection; this protein interferes with activation of the NF-κB pathway, which is absent from plants (Kim et al., 2005). Therefore, we examined whether OspF and OspG have virulence activity in plants by introducing them heterologously into a phytopathogen, *Pst*, and monitoring its pathogenicity. First, we used an AvrRpt2-derived T3SS reporter system (Figure S5A; (Mudgett, 2005). AvrRpt2^101-255^ is sufficient to induce cell death but lack in delivery to the plant cells. When *Arabidopsis* leaves were syringe-infiltrated with various *Pst* strains which express effectors fused to AvrRpt2^101–255^, we observed that *Pst* producing OspF:AvrRpt2^101–255^ or OspG:AvrRpt2^101–255^ successfully induced cell death at 1 day after delivery implying that both OspF and OspG are successfully delivered into *Arabidopsis* cells via *Pst* T3SS (Figure 5A). Next, we examined whether the virulence of *Pst* was increased by expression of *Shigella* OspF or OspG. Production of OspF:HA and OspG:HA by *Pst* was confirmed by immunoblotting with an anti-HA antibody (Figure S5B, S5C). Plants infected with *Pst* cells producing OspF: HA or OspG: HA showed more severe symptoms than plants infected with the empty vector control (Figure 5B). In addition, the number of bacterial cells was 10 times higher than in the plants infected with the empty vector control (Figure 5C). Taken together, these results indicate that OspF and OspG preserve their function as virulence proteins in plant hosts.

**Figure 5.**
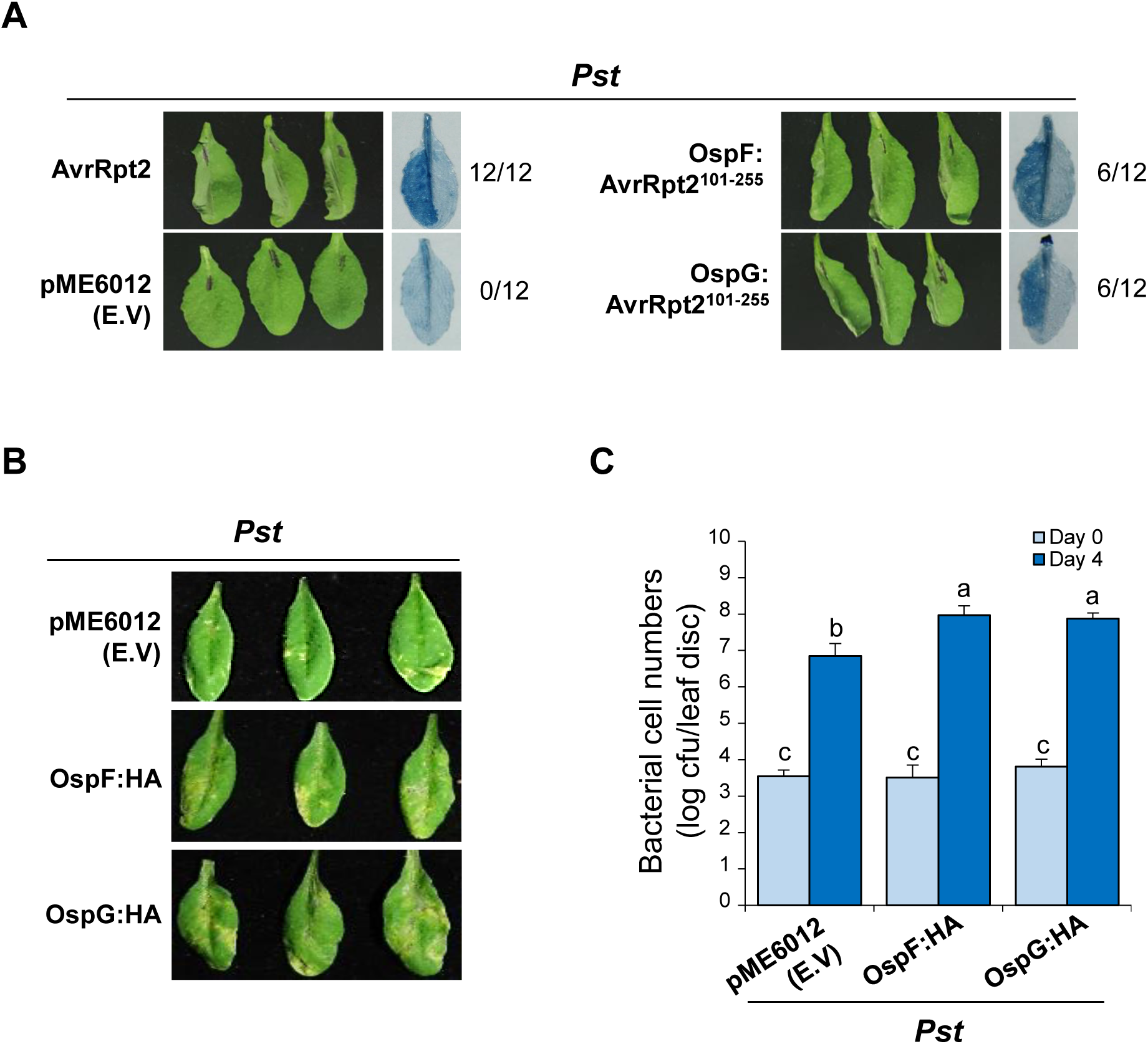
Increased virulence of phytopathogen *Pst* in the presence of *Shigella* effectors OspF or OspG. (A) *Pst* strains expressing the *Shigella* effector proteins were used in *Arabidopsis* for effector secretion assay. *Arabidopsis* leaves were syringe-infiltrated with *Pst* cells (5 × 10^7^ cfu/ml) producing AvrRpt2, carrying the empty vector (pME6012), and producing OspF:AvrRpt2^101–255^ or OspG:AvrRpt2^101–255^. The hypersensitive response in the infiltrated leaves was monitored using the trypan blue staining and photographed at 1 dpi. (B, C) *Arabidopsis* plants were sprayed with *Pst* cells carrying an empty vector (pME6012), or with cells producing OspF:HA or OspG:HA (5 × 10^7^ cfu/ml). (B) Disease symptoms were monitored for 4 d after spraying. (C) Proliferation of *Pst* cells producing *Shigella* effector proteins in *Arabidopsis* at 0 and 4 d after spray-inoculation. Bars represent the mean ± SD of six replicates, and different letters indicate significant differences between samples (*P* < 0.05, two-way ANOVA). All experiments were repeated at least three times.

On the other hand, *in vitro* studies demonstrated that OspF exerts a phosphothreonine lyase activity and irreversibly removes phosphate groups from MAPK (Arbibe et al., 2007; Li et al., 2007). To investigate whether the virulence associated with *Shigella* OspF in plants was linked to the same mechanism of action as in animals, we used a phenotypic screening system involving a MEK2 (a tobacco MAP kinase kinase2) mutant, MEK2^DD^ (Yang et al., 2001; Kim and Zhang, 2004). MEK2^DD^, a constitutively active mutant of MEK2, induces cell death when overproduced in plant leaves In this experiment, an HA:MEK2^DD^ clone fused to an HA epitope tag at the N-terminus was used to monitor expression of MEK2^DD^. As expected, co-production of HA:MEK2^DD^ and GFP (control) resulted in pronounced cell death (Figure S6A). Co-production of OspF:GFP, but not OspG:GFP, and MEK2^DD^ completely inhibited the MEK2^DD^-induced cell death (Figure S6A). The production of the two effectors fused with GFP was then evaluated in the MEK2^DD^-producing plant leaves; the production of OspF:GFP was apparent, but that of OspG:GFP was not (Figure S6B), even though both proteins were stably produced in the absence of MEK2^DD^ (Figure S6B). The degradation of the OspG protein might be associated with the activation of MAPK. Indeed, as assessed by immunoblotting with specific anti-phosphorylated MAPK antibodies, MAPK phosphorylation was very weak in the OspF:GFP-producing plant samples in comparison with GFP- or OspG:GFP-producing plant samples (Figure S6B). The production of MEK2^DD^ protein was confirmed in all samples using anti-HA antibodies (Figure S6B). These observations strongly suggest that the *Shigella* effector OspF inhibits plant immune responses by inhibiting activation of MAPK (as in humans), and that OspG induces immunosuppression in plants by targeting distinct MAPK pathways.

### OspF or OspG affects *Shigella* proliferation in plants

To confirm the role of the OspF or OspG proteins in the interaction between *Shigella* and plants (as in human cells), we inoculated plants with *S. f* 5a mutants lacking the OspF or OspG proteins and examined their growth. The growth of mutants lacking *ospF* or *ospG* was as deficient as that of the virulence plasmid-deficient mutant BS176 (Figure 6A). Reduced growth of *S. f* 5a *ΔospF* or *S. f* 5a *ΔospG* mutants was entirely restored by complementation of the mutation, indicating that these two effector proteins are indeed important for bacterial growth in plants (Figure 6A). Next, we monitored the activation of MAPKs to determine whether plant immune suppression was affected by the deletion of *ospF* or *ospG*. As shown in Figure 6B, *S. f.* 5a Δ*ospF* induced stronger phosphorylation of MAPKs than wild-type *S. f* 5a; this was offset by complementation with OspF. By contrast, phosphorylation of MAPK in *S. f* 5a *ΔospG-*inoculated plants was no different from that in plants inoculated with wild-type *S. f* 5a (Figure 6B). These results are consistent with the inhibition of MAPK-induced immune responses by OspF, but not by OspG, using the plant expression binary vector described above (Figure S6). Taken together, these results indicate that *Shigella* effectors OspF or OspG play an important role in increasing bacterial proliferation in both plant and animal hosts.

**Figure 6.**
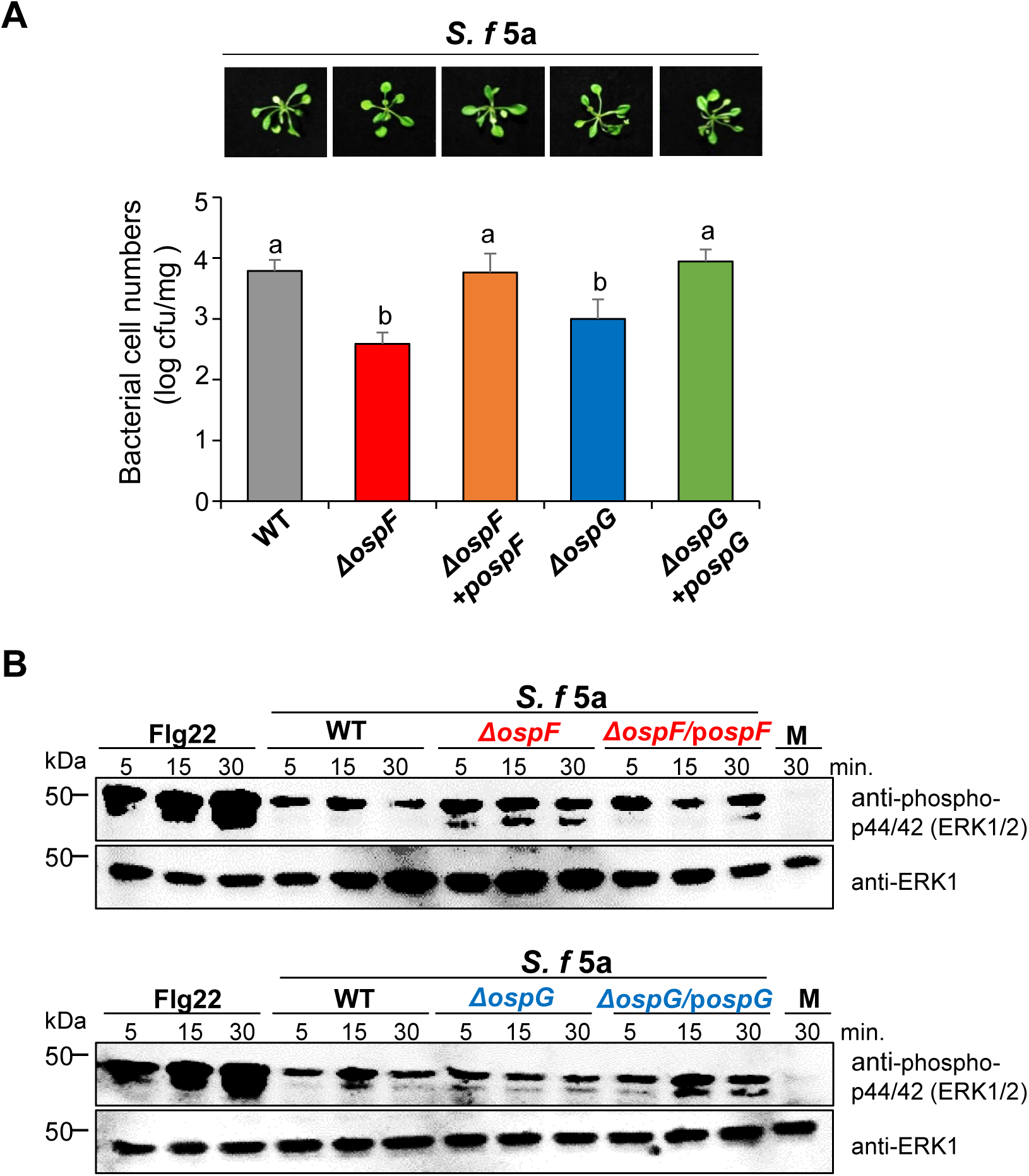
The *Shigella* effector proteins OspF and OspG underpin the bacterial virulence in plants. (A) The virulence of the OspF- or OspG-deficient *Shigella* mutants is impaired in the *Arabidopsis* model. *Arabidopsis* seedlings in 1/2 MS medium were flood-inoculated with *S. f* 5a, Δ*ospF*, Δ*ospF+*p*ospF*, Δ*ospG*, or Δ*ospG+*p*ospG* (5 × 10^5^ cfu/ml). Photographs of representative disease symptoms were taken, and bacterial cell numbers *in planta* were determined, at 3 dpi. The bars represent the mean ± SD of three replicates. The different letters indicate significant differences between samples (*P* < 0.05, one-way ANOVA). (B) MAPK activation in *Arabidopsis* plants in response to infection by ospF or ospG deletion mutants. *Arabidopsis* leaves were infiltrated with each bacterial strain as described previously and then collected at the indicated times. Anti-phospho-p44/42 (ERK1/2, upper panels) or anti-ERK1 (lower panels) antibodies were used for immunoblotting. All experiments were repeated at least three times, with similar results.

### Delivery of OspF and OspG into plant cells during *Shigella* inoculation

To visualize direct delivery of type III effectors to plant cells by *Shigella*, we took advantage of a newly developed split superfolder green fluorescent protein (sfGFP^OPT^) system (Park et al., 2017). In this system, the non-fluorescent GFP protein (GFP1-10) with β –strand 1 to 10 is expressed in the host cell and the bacterial effector is linked to the 11th β-strand (GFP11). Only when the effector fused with GFP11 is introduced into plant cells via bacterial T3SS, fluorescent signals are induced through GFP reconstitution (Park et al., 2017; Figure 7A). First, in order to select a host plant expressing GFP1-10 capable of exploring the delivery of OspF and OspG, we observed the subcellular localization of OspF:GFP and OspG:GFP by expressing them in *Nicotiana benthamiana* leaves using the *Agrobacterium* system (Figure S7). The fluorescence signal for both proteins was strong in the plant cell nucleus; OspF:GFP fluorescence was also observed along the cytoplasmic membrane, and punctate OspG:GFP fluorescence was observed in the cytosol (Figure S7C). However, immunoblot of OspG:GFP expression in *Nicotiana benthamiana* detected OspG:GFP as well as free GFP (Figure S7B), suggesting that overexpression of free GFP might be responsible for the cytosolic GFP expression. Based on these results, we choose *Arabidopsis* plants expressing sfGFP1-10 (sfGFP1-10^OPT^) in nucleus or cytosol and the OspF and OspG were fused with 11th β-strand of sfGFP, respectively (Figure S8A). GFP signals were observed at nucleus in both *S. f 5a*::OspF-sfGFP11 and *S. f 5a*::OspG-sfGFP11 flood-inoculated *Arabidopsis* leaves at 3 hpi, respectively, while control *S. f* 5a WT did not (Figure 7B). In contrast, when *Arabidopsis* seedlings expressing cytosolic sfGFP1-10^OPT^ were flood inoculated with *S. f* 5a producing OspF-sfGFP11 or OspG-sfGFP11, we detected no reconstituted GFP signals in all tested bacteria (Figure S8B). This suggests that *Shigella* effectors OspF and OspG were successfully secreted into *Arabidopsis* cells through the *Shigella* T3SS. In the host human cell, OspF localization is nuclear (Zurawski et al., 2006; Arbibe et al., 2007) and that of OspG is nuclear and cytoplasmic (Kim et al., 2005; Zhou et al., 2013; de Jong and Alto, 2014), confirming that the subcellular localization in plant cells is similar to that of animal cells. Taken together, the *Shigella* effectors OspF and OspG can be delivered into plant cells through T3SS.

**Figure 7.**
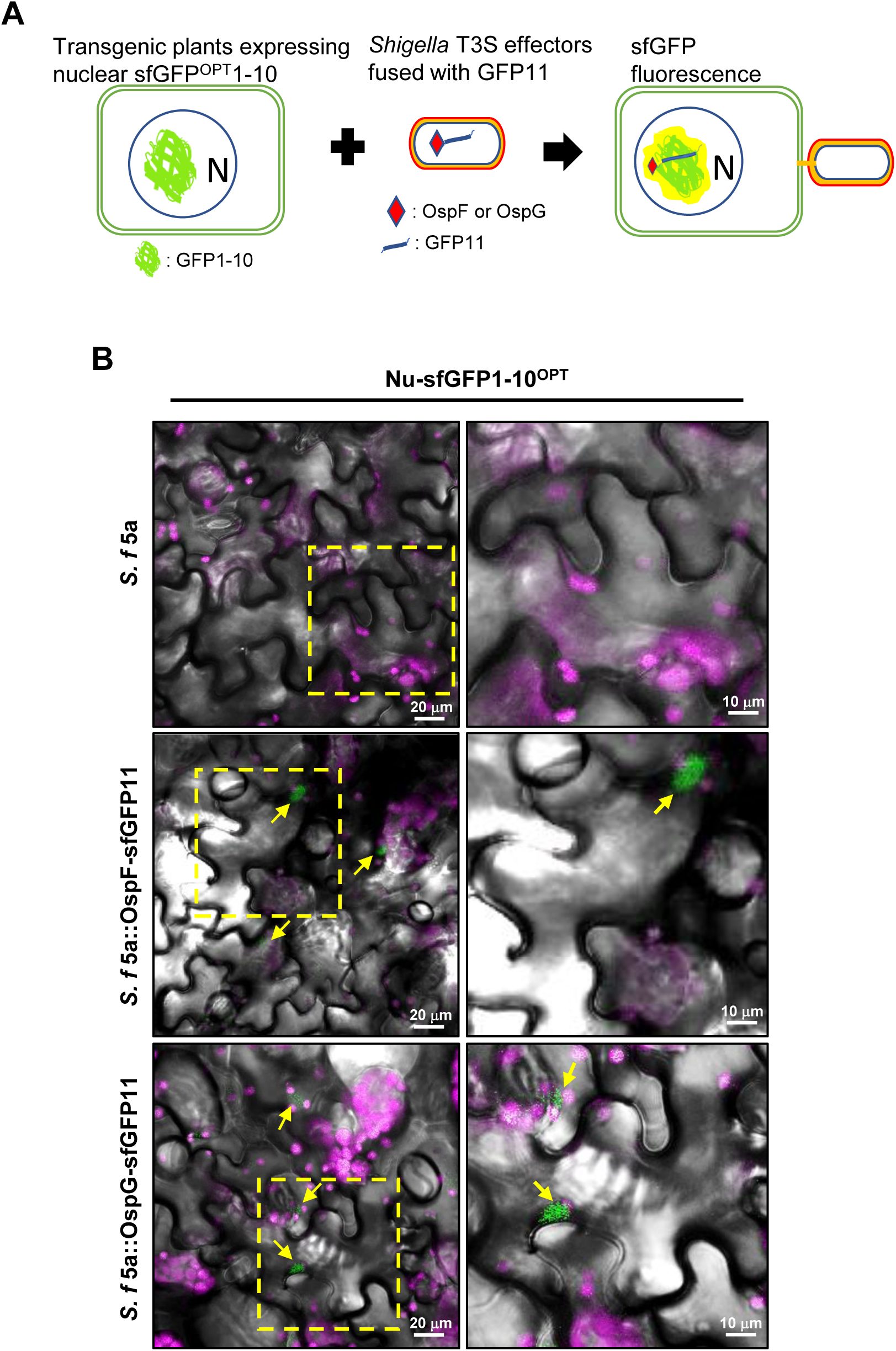
*Shigella* T3S effectors OspF and OspG with sfGFP11 are delivered through *S. flexneri* 5a into *Arabidopsis.* (A) Illustration of a split sfGFP^OPT^ system for monitoring the transfer of bacterial effectors by T3SS to the nucleus of host cells. (B) Transgenic *Arabidopsis* seedlings expressing sfGFP1-10 in 1/2 MS medium were flood-inoculated with *S. f* 5a containing OspF-sfGFP11 or OspG-sfGFP11(5 × 10 cfu/ml). 3 hours after inoculation reconstituted sfGFP signals in the nucleus of *Arabidopsis* leaf epidermal cells were observed under a fluorescence confocal microscope. Yellow arrows indicate the complemented sfGFP signals in nuclei. Yellow perporated-boxed regions were magnified at the right column. The magenta represents auto-fluorescence of chlorophyll. Bar, 10 μm and 20 μm.

## Discussion

In the current study, we investigated the interaction of the human pathogenic bacterium, *Shigella*, with an alternative host, the *Arabidopsis* plant. We demonstrated that four *Shigella* strains, *S. b*, *S. s*, *S. f* 2a, and *S. f* 5a, invade and colonize *Arabidopsis* to different extents. Different symptoms and bacterial proliferation rates for different *Shigella* strain suggested that the variability of the plant interaction mechanisms among the strains, e.g., the adherence and multiplication, might contribute to the differences. Differences in the nutritional requirements of bacterial strains may constitute another reason for the differences in the growth rates within the *Arabidopsis* host.

It has been demonstrated that, in human, *Shigella* initially enters the epithelial layer via the M cells through transcytosis, leading to the invasion of the basolateral surfaces of the intestinal epithelial cells. A subsequent gut inflammation leads to the activation of the innate immune response (Phalipon and Sansonetti, 2007). We demonstrated, in addition to the human host, *Shigella* invades *Arabidopsis* through vascular tissues and leaf stomata pores (Figure 2-4). In particular, *S. s* and *S. b* formed relatively wide clusters in the surrounding areas, including the guard cells (Figure 2), similar to *Pseudomonas syringae* which has built a host-pathogen relationship with Arabidopsis (Panchal et al., 2016). Interestingly, we found that the four studied-strains of *Shigella* associate with the plant cells and induce different plant responses. The bacterial loads of *S. s in planta* were relatively higher than those of the other strains. By contrast, inoculation of *S. f* 5a was associated with lower bacterial proliferation and less severe symptoms than observed for other strains (Figure 1-3). *S. f* 2a and *S. f* 5a, which belong to the same serogroup (Lindberg et al., 1991), elicited distinctly different plant responses with respect to disease symptoms, suggesting that the virulent effectors may play a relatively more important role in *Shigella*-plant interactions than PAMPs. These observations also indicate that specific plant immune systems may be useful in the search for novel virulence factors expressed by different *Shigella* strains.

Many Gram-negative bacterial pathogens utilize common infection strategies to colonize and invade plant and animal cells, and pathogenicity appears to depend on highly conserved T3SSs, which deliver the effector proteins to host cells (Buttner and Bonas, 2003). By using avirulent mutant strains, we were able to show that effectors that regulate the pathogenesis of shigellosis in humans also play a central role in regulating interactions with *Arabidopsis*. We showed that secretion of T3S effectors is required for the *Shigella* proliferation and attachment in plants (Figure 4, Figure S4). Furthermore, the effector proteins impacted MAPK*-*dependent/independent immune responses in *Arabidopsis* (Figure 4, 6). These observations further support the suggestion that the function of the main effector proteins of *Shigella* appears to be conserved in plant and animal hosts, and that this contributes to bacterial intracellular survival or suppression of the host defense against the pathogens. Reduced colonization of T3SS-deficient pathogenic *E. coli* in plants was previously reported, they suggested that *E. coli* uses the T3SS apparatus for attachment to leaves, rather than for bacterial growth inside plants (Shaw et al., 2008). The relevance of T3SS for multiplication of *Salmonella* in plants remains unclear due to the different effects of T3SS function on *Salmonella*-plant interactions (Iniguez et al., 2005; Schikora et al., 2011; Shirron and Yaron, 2011).

In general, animal bacterial pathogens with short needle lengths have been thought to be difficult to secrete T3S effectors on plant cells due to their cell walls and have been regarded as a major cause of not infecting plants. Recently, however, several studies have reported that the length and width of bacterial T3SS depend on the type of host cell, the environment, and the infection cycle of the bacteria (Deane et al., 2010). In this paper, we showed direct evidence that the effector proteins of *Shigella* were delivered into plant cells through T3SS utilizing recently developed the split GFP technology (Figure 7). Further study, identifying plant factors that trigger and control *Shigella*’s T3SS assembly will be a great help in developing a strategy to prevent the spread of *Shigella* through plants.

Expression of the T3SS of *Shigella* is regulated at the transcriptional level and is activated at a permissive temperature (≥ 32°C) (Tobe et al., 1991; Campbell-Valois and Pontier, 2016). We were able to observe the expression of the T3SS genes of *Shigella* under temperatures at which plants grow (22 ± 3°C) (Figure 5A). A recent study showed that elevation of the temperature increases T3SS-mediated virulence of the phytopathogen *Pst* in plants, which is in contrast with the negative effect of high temperature on expression of the T3SS of *Pst in vitro* (Huot et al., 2017). Regardless of the temperature of host cells, it will be interesting to determine whether *Shigella* regulates T3SS gene expression *in vivo* and to identify factors that influence T3SS gene expression other than plant temperature.

*Salmonella* strains capable of proliferating on plant leaves and actively entering plant tissues, root hairs, or trichomes were recently shown to exhibit virulence in animals (Barak et al., 2011; Golberg et al., 2011; Schikora et al., 2011). We demonstrated that *Shigella* strains actively colonize the surface of and inside *Arabidopsis* leaves and root tissues (Figure 1–3) and that bacteria recovered from plants maintain expression of pathogenic proteins (Figure S3). Collectively, these findings suggest that, similar to *Salmonella*, *Shigella*-inoculated plants are a serious risk to food safety and that contamination of plants is another route underlying infection of *Shigella*, an important human pathogenic bacterium. In this study, we only observed plants artificially inoculated with *Shigella* in a laboratory environment. Therefore, to confirm the food safety concern surrounding *Shigella*-inoculated plants, the ability of *Shigella* to infect a variety of crops grown in the field should be tested. The pathogenicity of plant-isolated *Shigella* in animals should also be investigated.

The current study provides new insights into host invasion mechanisms utilized by *Shigella* to interact with an alternative host, the plant *Arabidopsis*. Studying trans-kingdom pathogenesis involving human-adapted pathogens, such as *Shigella*, may uncover novel pathogenic mechanisms uniquely activated in response to specific hosts. When we isolated the two *Shigella* effectors OspF and OspG, and produced them in plant cells, their localization coincided with that in the animal cells (Figure 7 and S6C), and it was apparent that the production of both proteins increased the virulence of plant pathogens (Figure 6). In addition, we confirmed that OspF inhibits the innate immune response of plants via the same enzymatic activity as in animals (Figure 5 and S6). In animals, OspG inhibits the host immune response by inhibiting the activity of NF-κB by blocking degradation of IκB (Ashida et al., 2015). Plants possess an IκB-like protein called NIM1 (Ryals et al., 1997); however, no other published studies have investigated whether an NF-κB–induced immune response exists in plants. In the current study, we demonstrated the ability of OspG to increase the phytopathogenicity of non-*Shigella* bacteria (Figure 5), and also observed degradation of OspG:GFP upon constitutive activation of MAPK signaling (Figure S6B). The existence of a plant immune signaling pathway similar to that of animal NF-κB, which would also be the target of OspG, may hence be assumed. Characterization of the previously unrecognized stress-activated mediators of the innate immunity in plants upon infection with food-borne pathogens, would help us to define the defensive functions of plants. Finally, the characterization of plants as an alternative host for food-borne pathogens will be critical in developing effective means to prevent their transmission and disease.

## Acknowledgments

We thank Dr. Myung Hee Kim for initial support with the preparation of pathogenic *Shigella* strains, Dr. Jean T. Greenberg for providing the pBAV178, pBAV179, and pME6012 plasmids, and Dr. Cha Young Kim for providing the HA:MEK2^DD^ clone. This work was supported by the KRIBB Initiative Program and the Basic Research Program of National Research Foundation of Korea grant (NRF-2017R1A2B4012820 to JMP and NRF-2018R1A5A1023599 to DC) funded by the Korea government (MSIT).

## Author contributions

JMP, SHJ, and JL conceived and designed the study. SHJ, JL, and DHL carried out the experiments. JMP, SHJ, JL, DWK, and EP analyzed the data. JMP, SHJ, JL, and EP wrote the manuscript. JMP, SHJ, EP, CMR and DC edited and discussed the manuscript and all authors agreed with the final version.

## Supporting Information

Additional Supporting Information may be found online in the Supporting Information tab for this article:

**Fig. S1** The growth of *Shigella* strains producing GFP or virulence-deficient mutants is not affected *in vitro* growth.

**Fig. S2** Comparison of Bacterial proliferation in *Arabidopsis* inoculated with *Shigella* and GFP-labeled *Shigella* by flooding

**Fig. S3** Expression of effector proteins of *Shigella* recovered from inoculated plants.

**Fig. S4** Spontaneous type III secretion system-deficient mutant of *S. sonnei* is reduced growth and symptoms in *Arabidopsis*.

**Fig. S5** Expression of OspF or OspG in *Pst* for virulence assay.

**Fig. S6** Co-expression of OspF:GFP suppresses tobacco MEK2^DD^-triggered cell death.

**Fig. S7** Expression GFP fused OspF or OspG in plant.

**Figure S8**. Delivery of OspF or OspG fused with GFP11 through T3SS

**Table S1** Bacterial strains and plasmids used in the study.

**Table S2** Sequences of PCR primers used for Gateway cloning.

**Table. S3.** PCR primer list and sequences used for qRT-PCR.

