## Supplemental information for "A human pathogenic bacterium *Shigella* proliferates in the plant through the adoption of type III effectors for Shigellosis"

**Table. S1. Bacterial strains and plasmids used in this study.**

| Bacterial strains / plasmids | References |
|---|---|
| *Shigella boydii* ATCC9210 | Obtained from ATCC |
| *Shigella sonnei* 482279 | (Holt et al., 2012) |
| *Shigella flexneri* 2a 2457T | (Wei et al., 2003) |
| *Shigella flexneri* 5a M90T | (Onodera et al., 2012) |
| *Pseudomonas syringe* pv. tomato DC3000 | (Buell et al., 2003) |
| *Pseudomonas syringe* pv. tomato DC3000 *hrcC*$^-$ | (Hauck et al., 2003) |
| *Escherichia coli* DH5α | (Hanahan, 1985) |
| *S. flexneri* 2a 2457T ΔVP | (Shim et al., 2007) |
| *S. flexneri* 5a BS176 | (Sansonetti et al., 1982; Wenneras et al., 2000) |
| *Shigella flexneri* 5a M90T Δ*ospF* | (Arbibe et al., 2007) |
| *Shigella flexneri* 5a M90T Δ*ospF* + p*ospF* | (Arbibe et al., 2007) |
| *Shigella flexneri* 5a M90T Δ*ospG* | (Kim et al., 2005) |
| *Shigella flexneri* 5a M90T Δ*ospG* + p*ospG* | (Kim et al., 2005) |
| pDSK-GFPuv | (Wang et al., 2007) |
| pME6012 | (Vinatzer et al., 2005) |
| pBAV178 | (Vinatzer et al., 2005) |
| pBAV179 | (Vinatzer et al., 2005) |
| pK7FWG2 | (Bhaskar et al., 2009) |
| pEP119T | (Park et al., 2017) |

**Table. S2. Sequences of PCR primer used for Gateway cloning.**

| Primer list | Nucleotide sequence |
|---|---|
| *attB ospF- F* | 5′-aaagcaggctTCAATATATCTATTTTATAG-3′ |
| *attB ospF- R* | 5′-gaaagctgggtcCTCTATCATCAAACGATA-3′ |
| *attB ospG- F* | 5′-aaagcaggctTCGATTTTTAAATATCGTAA-3′ |
| *attB ospG- R* | 5′-gaaagctgggtcTAAATATTTCCTGTTTAA-3′ |

**Table. S3. PCR primer list and sequences used for qRT-PCR.**

| Primer list | Nucleotide sequence |
|---|---|
| *ipaB- F* | 5′-TTGGGCGTCGACTCGAAAA-3′ |
| *ipaB- R* | 5′-ACTGCTGCAACTAGGACAAGAG-3′ |
| *ipaC- F* | 5′-CCTCACCACAAACTAACTCTAGCA-3′ |
| *ipaC- R* | 5′-GAGAAGTTTTATGTTCAGTTGACAGGGATA-3′ |
| *icsA- F* | 5′-CTCTCTGTAATCAATAAGGGCACGTT-3′ |
| *icsA- R* | 5′-CCACCGTAGCCATCATAACCATAA-3′ |
| *icsB- F* | 5′-CATTCCGCCGGGATACCA-3′ |
| *icsB- R* | 5′-GGCATAACCCATTTAGGGCATACTA-3′ |
| *virB- F* | 5′-TCTCGCGCGAAAGTCACT-3′ |
| *virB- R* | 5′-GTTCTGACGCGATTGGAAATAGAGA-3′ |
| *virF- F* | 5′-TCTGAGGAGGAGGTTTCTATCGATT-3′ |
| *virF- R* | 5′-GAAACAGCTGATAAAAGGCAAGCT-3′ |
| *16S- F* | 5′-GCGGTTTGTTAAGTCAGATGTGAAA-3′ |
| *16S- R* | 5′-GACTCAAGCTTGCCAGTATCAGAT-3′ |

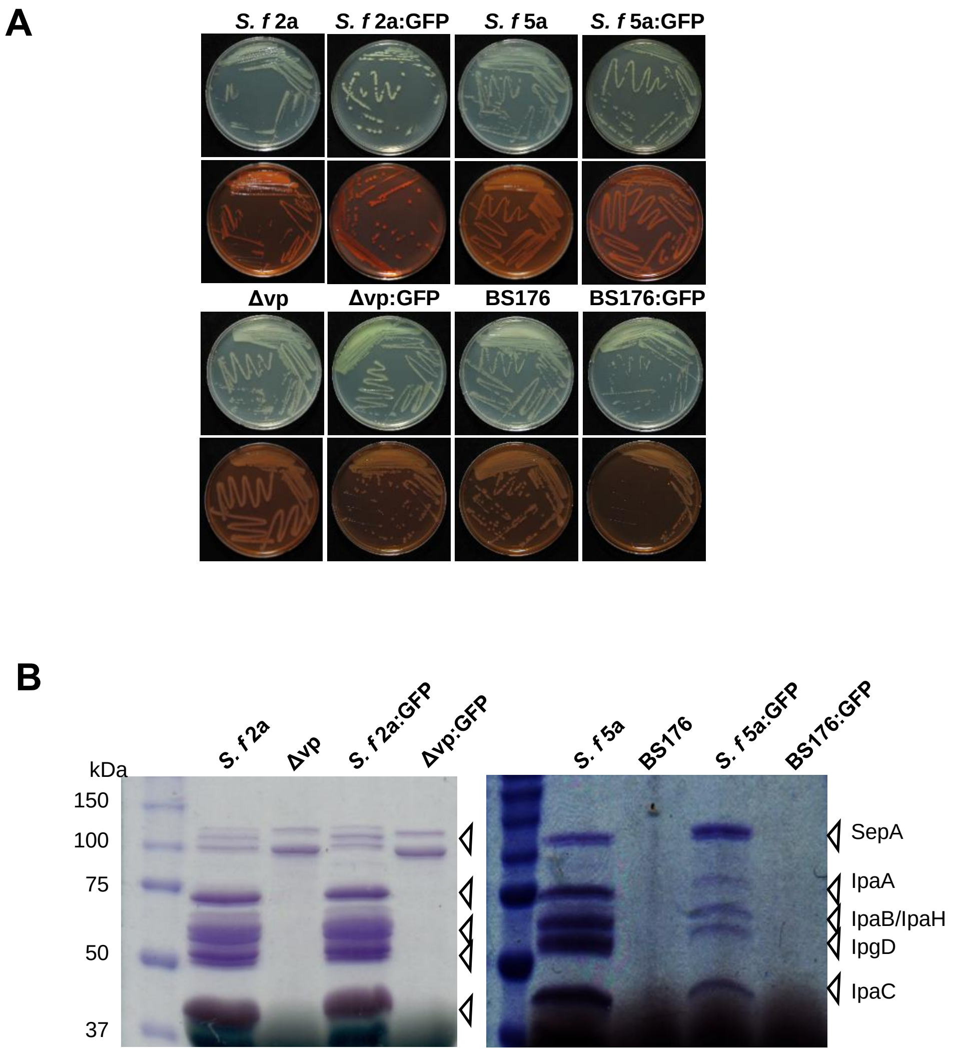


**Figure S1. The growth of *Shigella* strains producing GFP or virulence-deficient mutants is not affected *in vitro* growth.**

(A) *S. f* 2a, Δvp, *S. f* 5a, and BS176 showed normal growth phenotypes in the Luria-Bertani (LB) medium, but the avirulent mutant colonies were less colored on the tryptic soy agar medium containing Congo red dye. The photographs of representative plates were taken after 1 d of incubation at 37°C. (B) The *S. f* 2a, Δvp, *S. f* 5a, and BS176 cultures in the tryptic soy broth medium in presence of Congo red for 3 h at 37 °C were harvested to analyze the secreted effector proteins. The precipitated supernatants by centrifugation were analyzed by SDS-PAGE and Coomassie blue R solution staining. The molecular weight markers are indicated on the left of the Coomassie-stained gels, and the positions of the various proteins according to molecular weight are indicated to the right.

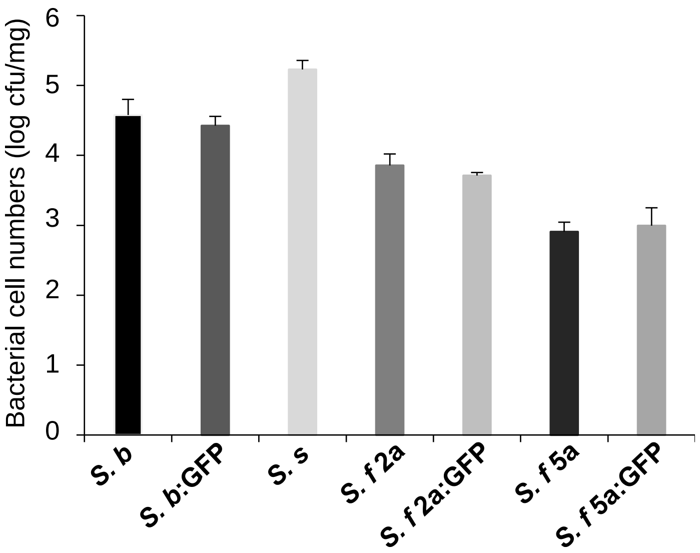


**Figure S2. Comparison of Bacterial proliferation in *Arabidopsis* inoculated with *Shigella* and GFP-labeled *Shigella* by flooding.**

*Arabidopsis* seedlings in 1/2 MS medium were flood-inoculated with *Shigella* suspensions ($5 \times 10^5$ cfu/ml). Bacterial cell numbers were evaluated on day 3 after the inoculation. The bars represent the means ± SD of three replicates. The experiments were repeated at least three times, and representative results are shown.

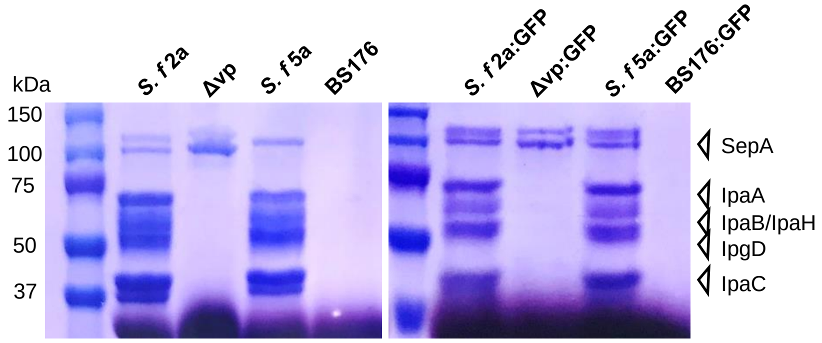


**Figure S3. Expression of effector proteins of *Shigella* recovered from inoculated plants.**

*Arabidopsis* grown *S. f* 2a, Δvp, *S. f* 5a, and BS176 were used to analyze the secreted effector proteins. *Shigella* were harvested from inoculated *Arabidopsis* at 3dpi and cultivated for secretion assay. The precipitated supernatants of cultivated cells by centrifugation were analyzed by SDS-PAGE and Coomassie blue R solution staining.

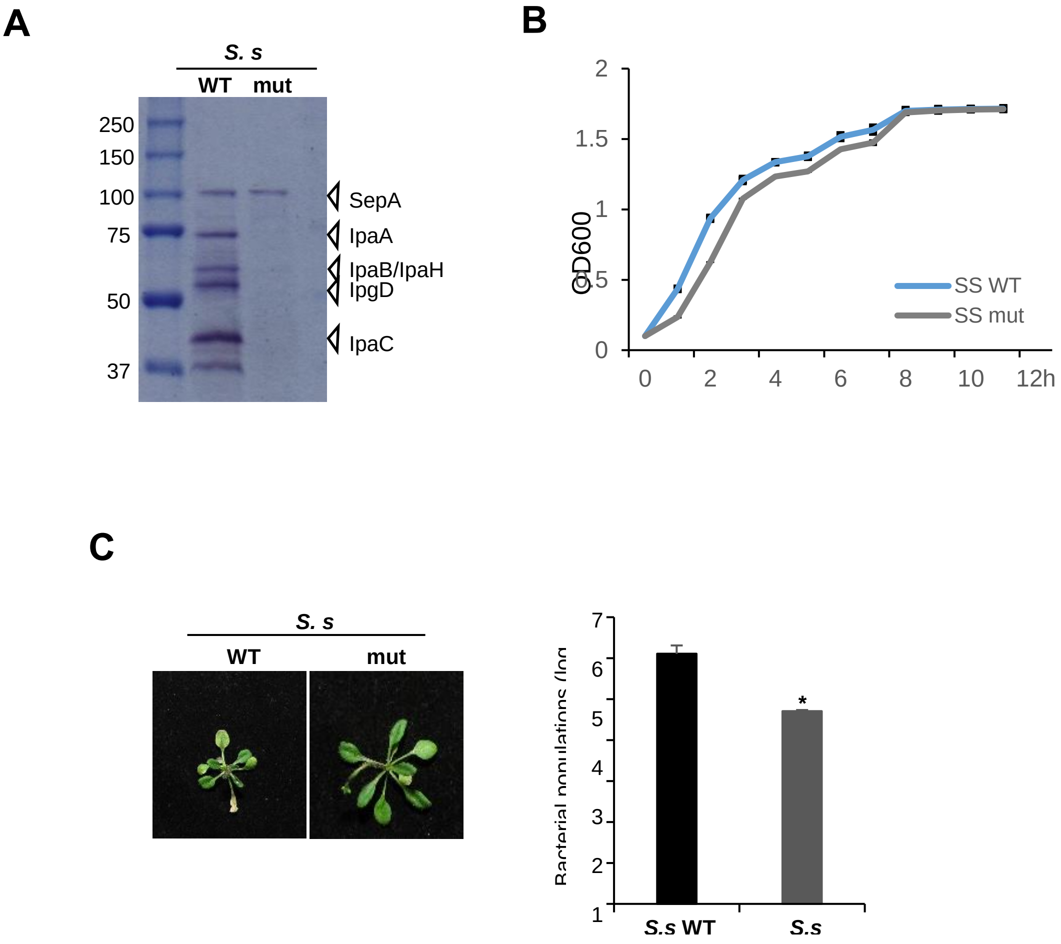


**Figure S4. Spontaneous type III secretion system-deficient mutant of *S. sonnei* is reduced growth and symptoms in *Arabidopsis*.**

(A) Cultures of the *S.s* (WT) and type III secretion system-deficient mutant of *S.s* (mut) were harvested in presence of Congo Red for 3 h at 37$^{o}$C to analyze the secreted effector proteins. Precipitated supernatants were analyzed by SDS PAGE and Coomassie stain. The molecular weight markers are indicated on the left of the Coomassie-stained gel, and the positions of the various proteins are indicated to the right. (B) Bacterial growth curve in LB media was identified in *S. s* WT, or mut. The bars represent the means ± SD of three replicates. (C) *Arabidopsis* seedlings in 1/2 MS medium were flood-inoculated with *S. s* WT, or mut ($5 \times 10^{5}$ cfu/ml). Photographs of representative disease symptoms were taken and bacterial populations in planta were measured, at 3 dpi. The error bars represent the SD of three replicates, and the asterisks indicate significant difference compared with *S. s* WT (* P<0.05, *t*-test).

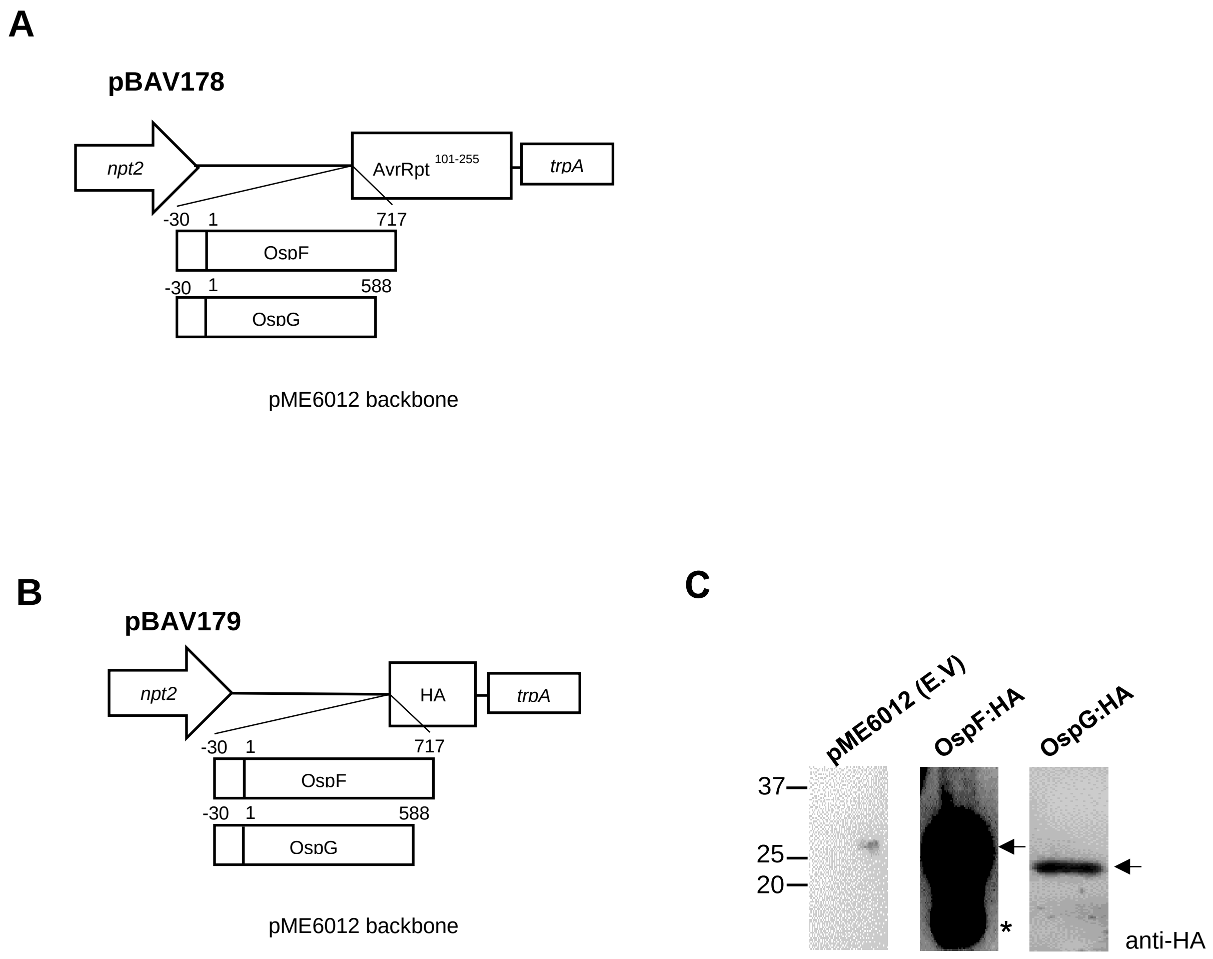


**Figure S5. Expression of OspF or OspG in *Pst* for virulence assay.**

(A) A schematic representation of pBAV178 vector. (B) A schematic representation of pBAV179 vector. (C) *Pst* strains expressing the *Shigella* effector proteins were used in *Arabidopsis* virulence spray assay. OspF or OspG proteins were identified using the anti-HA antibody by immunoblotting. Asterisk indicates the degraded form of the effector proteins.

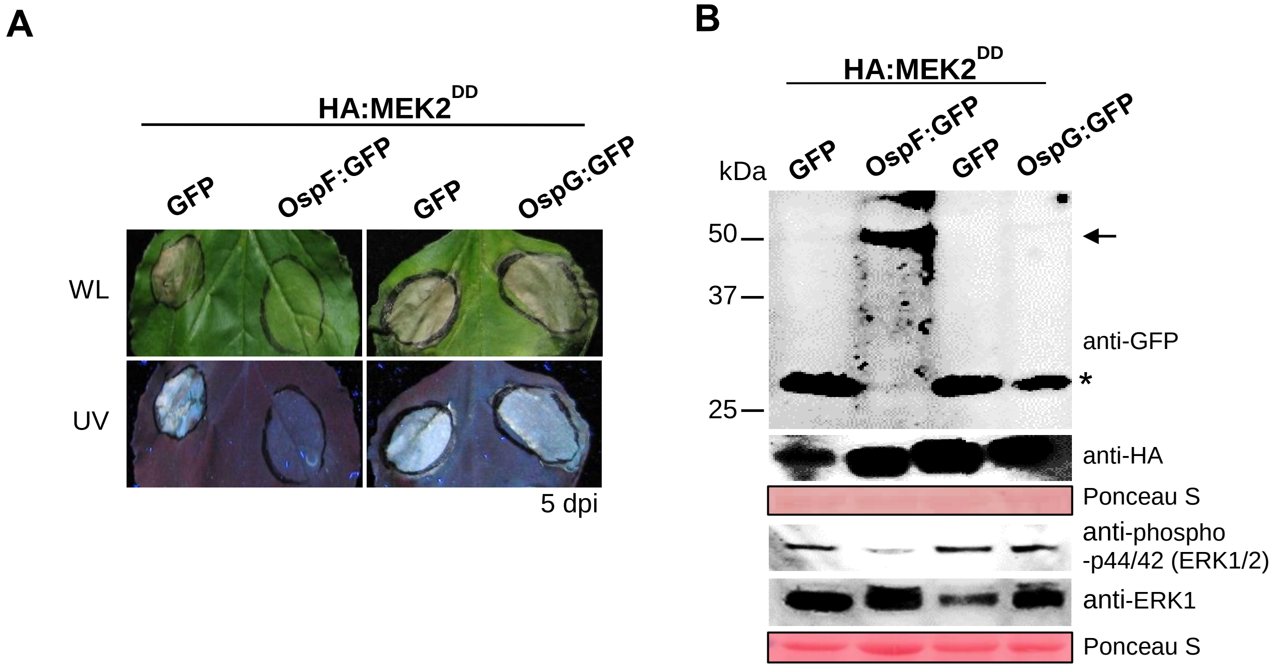


**Figure S6. Co-expression of OspF:GFP suppresses tobacco MEK2$^{DD}$-triggered cell death.**

(A, B) GFP, OspF:GFP, or OspG:GFP was produced with HA:MEK2$^{DD}$ in *N. benthamiana* leaves upon infiltration by *Agrobacterium* carrying the appropriate expression constructs ($OD_{600}$ 0.4). (A) Production of OspF:GFP suppressed MEK2$^{DD}$-dependent cell death by 5 d after the co-infiltration. (B) Production of specific proteins in samples from panel A was analyzed by immunoblotting using anti-GFP, anti-HA, anti-phospho-p44/42 (ERK1/2), and anti-ERK1 antibodies. Ponceau S was used to stain the RuBisCo protein (loading control). The asterisk indicates the size of the GFP protein not associated with effector proteins.

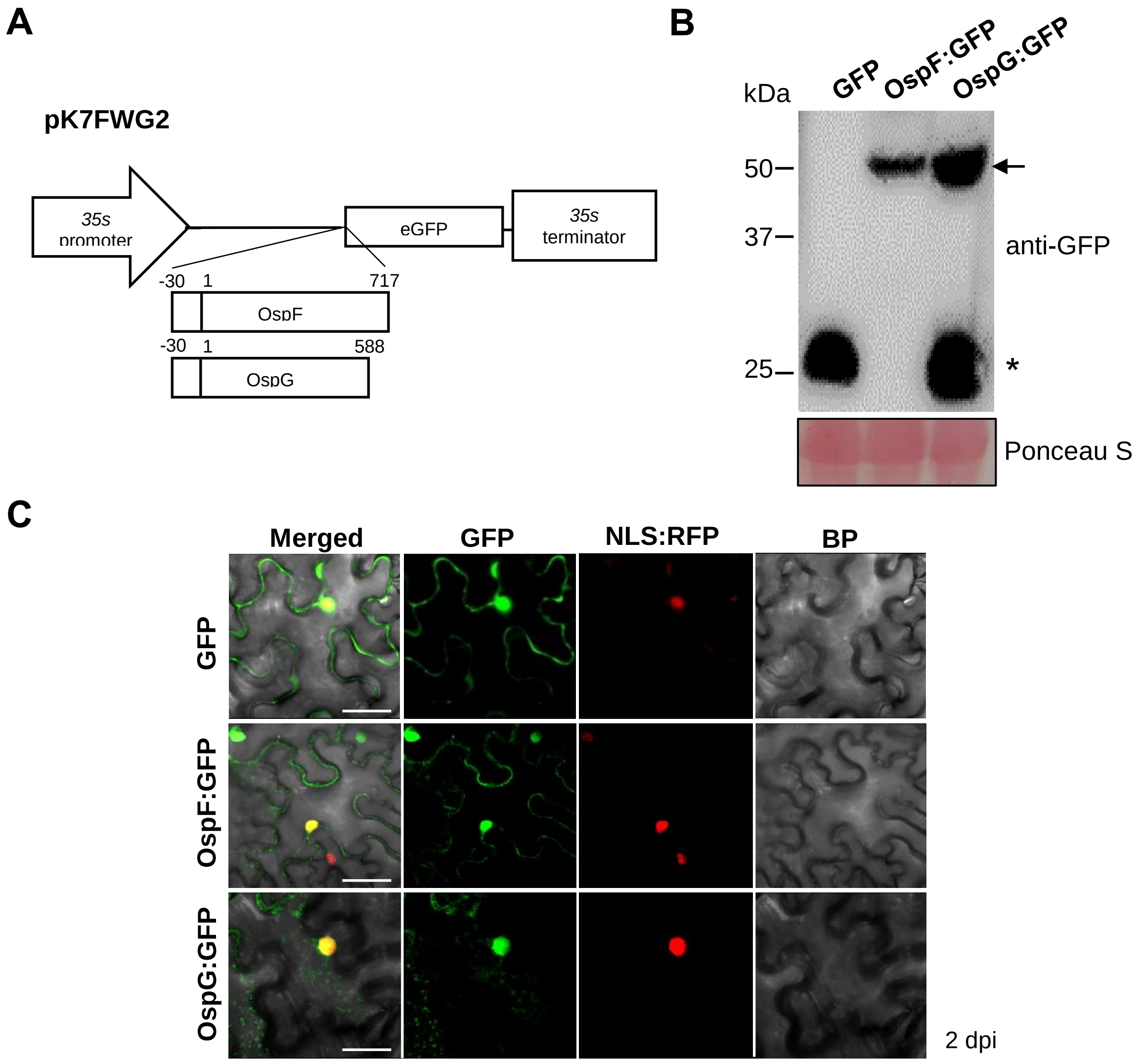


**Figure S7. Expression GFP fused OspF or OspG in plant.**

(A) A schematic representation of pK7FWG2 vector. (B) GFP, OspF:GFP, or OspG:GFP was produced in *N. benthamiana* leaves upon infiltration with *Agrobacterium* carrying the appropriate expression constructs ($OD_{600}$ 0.4). The production of the *Shigella* effector proteins in *N. benthamiana* 2 d after the infiltration was evaluated by immunoblotting using anti-GFP antibodies. The RuBisCo protein was stained by Ponceau S as a loading control. (C) Subcellular localization of GFP, OspF:GFP, OspG:GFP, and NLS:RFP (a nuclear localization marker) in *N. benthamiana* leaves. Bar, 50 µm.

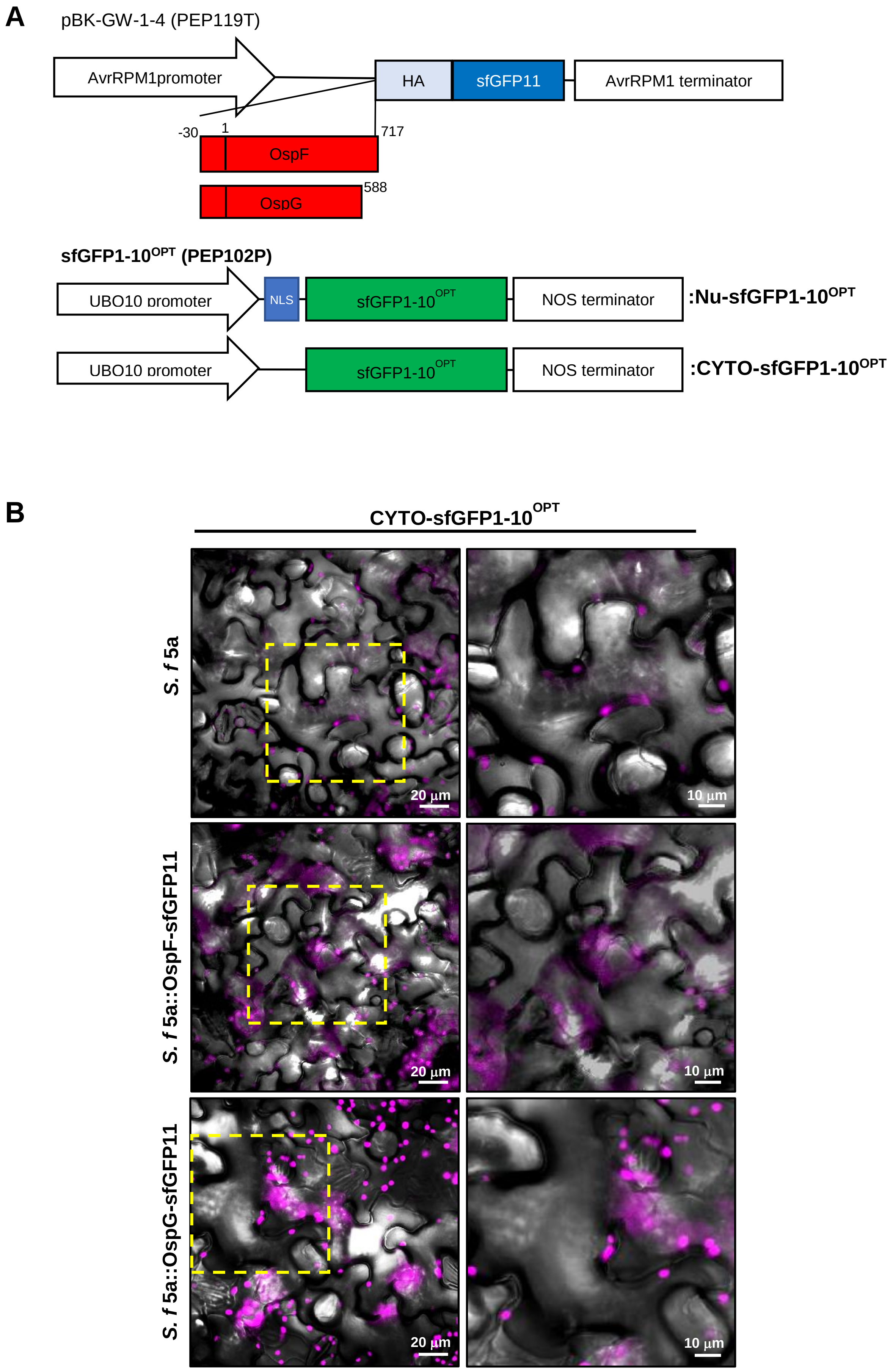
A
pBK-GW-1-4 (PEP119T)
AvrRPM1promoter
HA
sfGFP11
AvrRPM1 terminator
-30
1
717
OspF
OspG
588
sfGFP1-10OPT (PEP102P)
UBO10 promoter
NLS
sfGFP1-10OPT
NOS terminator
:Nu-sfGFP1-10OPT
UBO10 promoter
sfGFP1-10OPT
NOS terminator
:CYTO-sfGFP1-10OPT
B
CYTO-sfGFP1-10OPT
S. f 5a
S. f 5a::OspF-sfGFP11
S. f 5a::OspG-sfGFP11
20 µm
10 µm
20 µm
10 µm
20 µm
10 µm

**Figure S8. Delivery of OspF or OspG fused with GFP11 through T3SS**

(A) A schematic representation of sfGFP$^{OPT}$ vector. (B) Transgenic *Arabidopsis* seedlings expressing cytosolic sfGFP1-10$^{OPT}$ in 1/2 MS medium were flood-inoculated with *S. f* 5a containing OspF-sfGFP11 or OspG-sfGFP11 ($5 \times 10^5$ cfu/ml). 3 hours after inoculation no sfGFP signals in *Arabidopsis* leaf epidermal cells were observed under a fluorescence confocal microscope. The magenta represents auto-fluorescence of chlorophyll. Bar, 20 μm and 10 μm.

### Materials and Methods

#### Analyzing of secreted effector proteins

Single colonies of *S. f* 2a, Δvp, *S. f* 5a, and BS176 from Congo red tryptic soy agar plates were incubated overnight with shaking at 37 °C in LB medium. Followed by subculture (1:100 dilution), medium were grown at 37 °C up to $OD_{600}$ of 0.3-0.4. Congo red was added to a final concentration of 200 μg/ml and incubated for 2-3h at 37°C. Culture medium were harvested to analyze secreted effector proteins. After centrifugation and TCA precipitation, supernatants were analyzed by SDS-PAGE and Coomassie blue R solution staining (Reinhardt and Kolbe, 2014).

#### Immunoblotting

Total protein was extracted from *Pst* or *Agrobacterium*-infected leaves (from three plants) and used for immunoblot analysis as described in main text. Antibodies for hemagglutinin (HA) (Cat No: S2930; Clontech Laboratories, Mountain View, CA, USA), or GFP (Cat No: sc-9996, Santa Cruz) were used.

#### *Agrobacterium*-mediated transient gene expression

The coding region of *ospF* or *ospG* was also transferred to pK7FWG2 (obtained from Ghent University, Belgium) by LR recombination to produce the GFP-fused *Shigella* effectors OspF:GFP and OspG:GFP (Karimi et al., 2002).

*A. tumefaciens* strain GV2260 harboring the *GFP*, *OspF:GFP*, or *OspG:GFP* genes driven by the 35S promoter was prepared as described previously (Lee et al., 2013). The inoculum ($OD_{600}$ = 0.4) was infiltrated into 4-week-old *N. benthamiana* leaves using a 1 ml needleless syringe. To observe $MEK2^{DD}$-triggered cell death suppression by *Shigella* effectors, *Agrobacterium* ($OD_{600}$ = 0.4) expressing HA:$MEK2^{DD}$ was mixed with *Agrobacterium* containing *GFP*, *OspF:GFP*, or *OspG:GFP* at a 1:1 ratio and infiltrated into the leaves of *N. benthamiana*. Each experiment was repeated using at least three leaves from the plant, and each experiment included at least three different plants.
